# Metabolic functional redundancy of the CYP9A subfamily members leads to P450-mediated *lambda*-cyhalothrin resistance in *Cydia pomonella*

**DOI:** 10.1101/2022.07.22.501203

**Authors:** Pei-Rong Li, Yu Shi, Yu-Xi Liu, Wei Wang, Di Ju, Ying-Shi He, Yu-Yun Zhang, Xue-Qing Yang

## Abstract

**BACKGROUND:** The evolution of insect resistance to pesticides poses a continuing threat to sustainable pest management. While much is known about the molecular mechanisms that confer resistance in model insects and few agricultural pests, far less is known about fruit pests.

**RESULTS:** Here we found that functional redundancy and preference of metabolism by cytochrome P450 monooxygenases (P450s) genes in the *CYP9A* subfamily confer resistance to *lambda*-cyhalothrin in *Cydia pomonella*, a major invasive pest of pome fruit. A total of four CYP9A genes, including *CYP9A61*, *CYP9A120*, *CYP9A121*, and *CYP9A122*, were identified from *C. pomonella*. Among these, *CYP9A120*, *CYP9A121*, and *CYP9A122* were predominantly expressed in the midgut of larvae. The expression levels of these P450 genes were significantly induced by LD_10_ of *lambda*-cyhalothrin and were overexpressed in a field-evolved *lambda*-cyhalothrin resistant population. Knockdown of *CYP9A120* and *CYP9A121* by RNA-mediated interference (RNAi) increased the susceptibility of larvae to *lambda*-cyhalothrin. In *vitro* assays demonstrated that recombinant P450s expressed in Sf9 cells can metabolize *lambda*-cyhalothrin, but with functional redundancy and divergence through regioselectivity of metabolism. CYP9A121 preferred to convert *lambda*-cyhalothrin to 2′-hydroxy-*lambda*-cyhalothrin, whereas CYP9A122 only generated 4′-hydroxy metabolite of *lambda*-cyhalothrin. Although possesses a relatively low metabolic capability, CYP9A120 balanced catalytic competence to generate both 2′- and 4′-metabolites.

**CONCLUSION:** Collectively, these results reveal that metabolic functional redundancy of three members of the CYP9A subfamily leads to P450-mediated *lambda*-cyhalothrin resistance in *C. pomonella*, thus representing a potential adaptive evolutionary strategy during its worldwide expansion.

## 1. INTRODUCTION

The codling moth, *Cydia pomonella* (L.) (Lepidoptera: Tortricidae) is a major global pest of pome fruits ^1–3^ and also an invasive agricultural pest species in China.^4^ The larvae of this notorious pest penetrate the center of the fruit, which causes the abscission of fruits, making it unmarketable, with annual damage estimated to be tens of millions of dollars.^4, 5^ This pest is native to Mediterranean Europe, but currently it can be found on six continents and imposes severe damage on pome fruit production globally.^4^ In China, since the localized population was documented in Xinjiang in 1957 ^6^, *C. pomonella* increasingly widened its distribution and has spread to Gansu, Ningxia, Inner Mongolia, Heilongjiang, Jilin, and Liaoning Provinces in recent years, closing into the main apple production areas in the Loess Plateau and Bohai Bay, and causing a serious threat to the development of the apple industry in China.^1, 3^ However, the mechanisms of the global spread of *C. pomonella* are largely unclear, which limits the development of prevention and control strategies for this invasive pest.

In almost all pome fruit-producing countries, effects to control the *C. pomonella* have mainly relied on synthetic insecticides such as *lambda*-cyhalothrin, particularly widely used.^5, 7, 8^ However, with the extensive application of insecticides, field-evolved resistance has been widely documented in *C. pomonella* worldwide^1, 8–10^ aside potential multiple occurring side effects of pesticides on non-target organisms.^11^ The evolution of resistance to insecticides not only demonstrated the striking ability of this species to adapt to insecticides stress, which may explain its worldwide expansion, but also an ongoing challenge to sustainable management of this species.^4^ Understanding the mechanisms underlying resistance to insecticides is useful for resistance management and developing efficient pest management strategies.^4, 12^ At least two major mechanisms, including target-site insensitivity and metabolic enzyme-based resistance, are known to cause insecticide resistance in *C. pomonella*.^8^ Among these two mechanisms, metabolic resistance is a more common mechanism with the increased detoxification enzymes such as cytochrome P450 monooxygenases (P450s), glutathione *S*-transferases (GSTs), or transporters like ATP binding cassette (ABC) transporters.^13^ Recently, overexpression of GSTs and ABC genes was involved in insecticide resistance in *C. pomonella*.^8, 10^ Besides, the increased production of P450s has been linked to resistance to insecticides including *lambda*-cyhalothrin in field-evolved resistant populations from Spain, France, Greece, Turkey, and China.^1, 8^ However, the underlying molecular mechanisms of P450-mediated insecticide resistance remain largely unknown in *C. pomonella*.

P450s, encoded by the *CYP* genes, are made up of a remarkable superfamily of enzymes.^14^ Despite the substantial progress achieved in the molecular analysis and identification of several P450s associated with insecticide resistance in arthropods, the exact role of P450s in insecticide resistance remains largely unknown.^15^ P450 genes distributed to Clan 3, particularly in *CYP6B* and *CYP9A* subfamilies, are thought to play crucial roles in insecticide resistance in insects.^16^ The transcript of *CYP9A61*, the first P450s gene identified from *C. pomonella*, was induced by *lambda*-cyhalothrin, chlorpyrifos-ethyl, imidacloprid, carbaryl, and deltamethrin, suggesting a potential role of this gene in insecticide detoxification.^7, 17^ Resistance of *C. pomonella* to deltamethrin and azinphos-methyl has been linked to the overexpression of a single P450 gene, *CYP6B2*.^4^ Despite the importance of P450 genes of the *CYP9A* subfamily in insecticide resistance, the molecular basis for overexpression of *CYP9A* genes in *lambda*-cyhalothrin-resistant strains has never been elucidated in *C. pomonella*.

In this study, we identified *CYP9A* genes and investigated whether they were involved in *lambda*-cyhalothrin resistance in *C. pomonella.* The results of this study may not only help us better understand the molecular basis of resistance to insecticides and develop new strategies for resistance management but also be useful for elucidating the mechanisms for the global spread of *C. pomonella*.

## 2. MATERIALS AND METHODS

### 2.1 Insects

A susceptible strain of *C. pomonella* (SS) is reared in the laboratory without insecticides exposure for more than 50 generations.^10^ A field field-evolved *lambda*-cyhalothrin-resistant population (ZW_R) of *C. pomonella*^18^ was used in this study. This strain was reared in the laboratory under the same conditions as the SS strain.

### 2.2 Reagents and chemicals

*Lambda*-cyhalothrin (99.2%, CAS: 91465-08-6) was purchased from MedChemExpress company (Monmouth Junction, New Jersey). Four fluorescent P450 model probe substrates, including 7-benzyloxy-resorufin (BR), 7-benzyloxy-4-trifluoromethyl coumarin (BFC), 7-ethoxy-4-trifluoromethyl coumarin (EFC), and 7-benzyloxymethoxy-resorufin (BOMR), were supplied by Sigma-Aldrich (Saint Louis, MO) or Life Technologies. Reduced nicotinamide adenine dinucleotide phosphate (NADPH) and the components of the NADPH regeneration system used for *in*-*vitro* metabolism were purchased from Sigma-Aldrich Company (Saint Louis, MO). HPLC solvents were purchased from Fisher Scientific (Pittsburgh, Pennsylvania).

### 2.3 Identification, sub-cloning, and phylogenetic analysis of P450 genes

The CYP9A sequences of representative species in Noctuidae including *C. pomonella*, *H. armigera*, *Bombyx mori*, *Spodoptera exigua,* and *Mamestra brassicae* (Table S1) were downloaded from National Center for Biotechnology Information (NCBI) and were used as queries for searches of TBLASTN (e-value < 10^-5^) against the *C. pomonella* transcriptome database (SRX371333) by using the Blast v2.5.0+.^19^ Then we used Clustal X to filter out the repeat sequences.^20^ The obtained sequences were mapped back to the genome database of *C. pomonella* deposited in InsectBase (http://v2.insect-genome.com/Organism/224).

Total RNA was extracted by Eastep Super Total RNA Extraction Kit (Promega, Shanghai, China) based on the manufacturer’s instructions. The concentration and quality of the RNA samples were determined by NanoDrop 2000 (ThermoFisher Scientific, Waltham, MA). The first-strand cDNA was prepared by GoScript^TM^ Reverse Transcription Mix (Promega, Madison, WI) using 1 μg of extracted RNA of *C. pomonella*. Specific primers (Table 1) were designed by Primer Premier 5.0, and were then synthesized by Sangon Biotech Co. Ltd. (Shanghai, China). PCR was performed using PrimeSTAR Max DNA polymerase (Takara, Dalian) to amplify the predicted *CYP9A* genes under the following cycling conditions: 94 for 1 min, followed by 35 cycles of 94 for 30 s, 55 for 30 s and 72 for 30 s, and a final extension at 72 for 2 min. The PCR products were gel-purified using Biospin Gel Extraction Kits (Bioer Technology Co., Ltd., Hangzhou, China), sub-cloned into the pMD 19-T vector (TaKaRa, Dalian, China), transformed into *Escherichia coli* DH5α (TaKaRa, Dalian, China) and finally sequenced at Shanghai Sangon Biotech Co. Ltd. T. ORFfinder (https://www.ncbi.nlm.nih.gov/orffinder/) was used to predict the open reading frame (ORF) of sequenced CYP9A genes. ExPASy Proteomics Server (https://web.expasy.org/compute_pi/) was used to predict the molecular weight (Mw) and isoelectric point (pI). Multiple sequence alignment was carried out using the ClustalW.^20^ The similarity of the sequences was determined and nucleotide and deduced amino acid sequences were predicted by DNAMAN (LynnonBioSoft, CA, USA). The conserved regions of P450 proteins (FxxGxxxCxG/A, PxxFxPxRF, A/GGx D/ETT/S, ExxR, and WxxR) and the secondary structures were predicted using PSIPRED.^21^ The 3D structure of CYP9A120, CYP9A121 and CYP9A122 were predicted using the SWISS-MODEL (http://swissmodel.expasy.org). A phylogenetic tree was constructed by MEGA 7.0 using the neighbor-joining method and 1000 bootstrap repeats.^22^

**Table 1.**
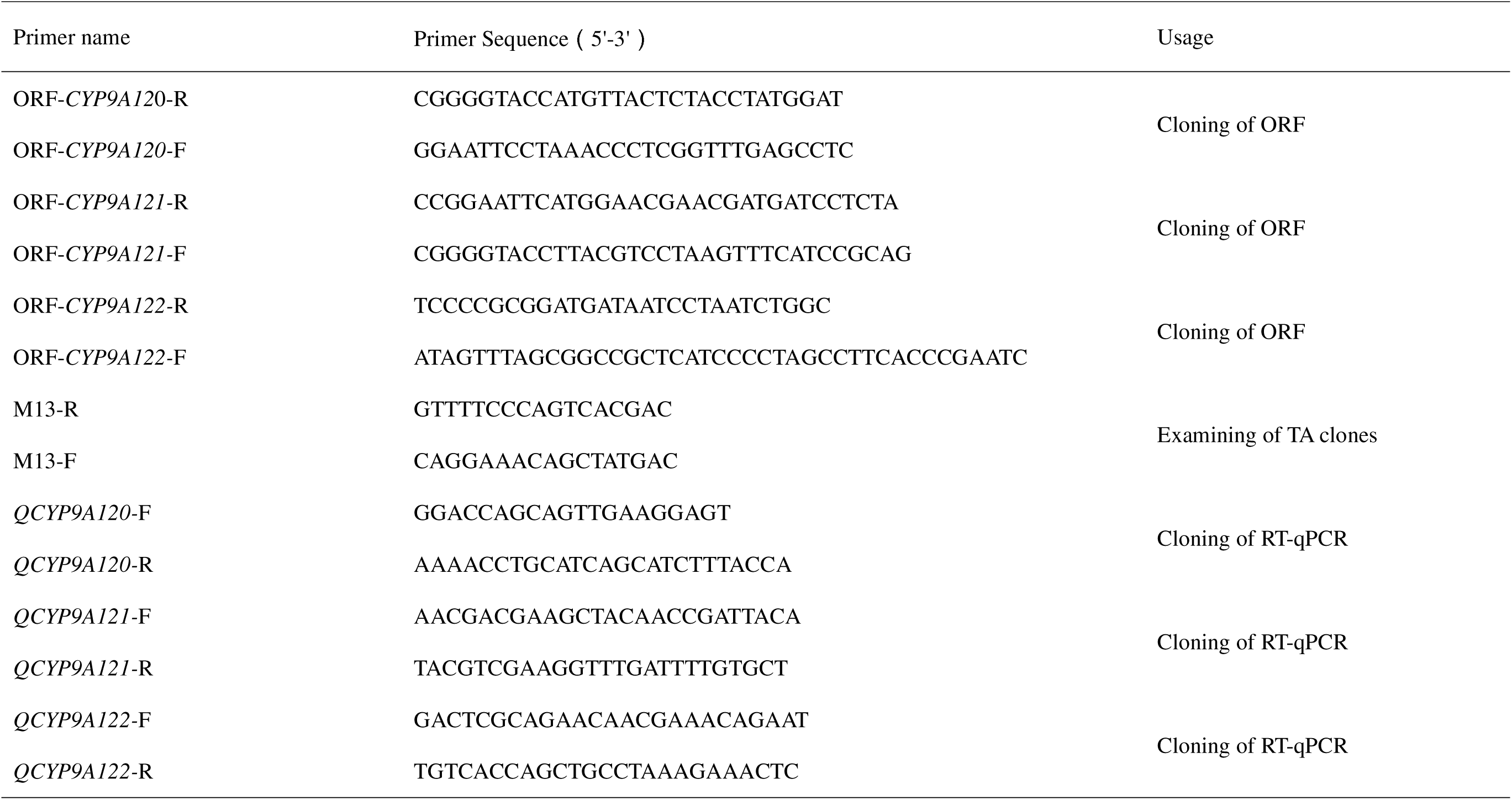
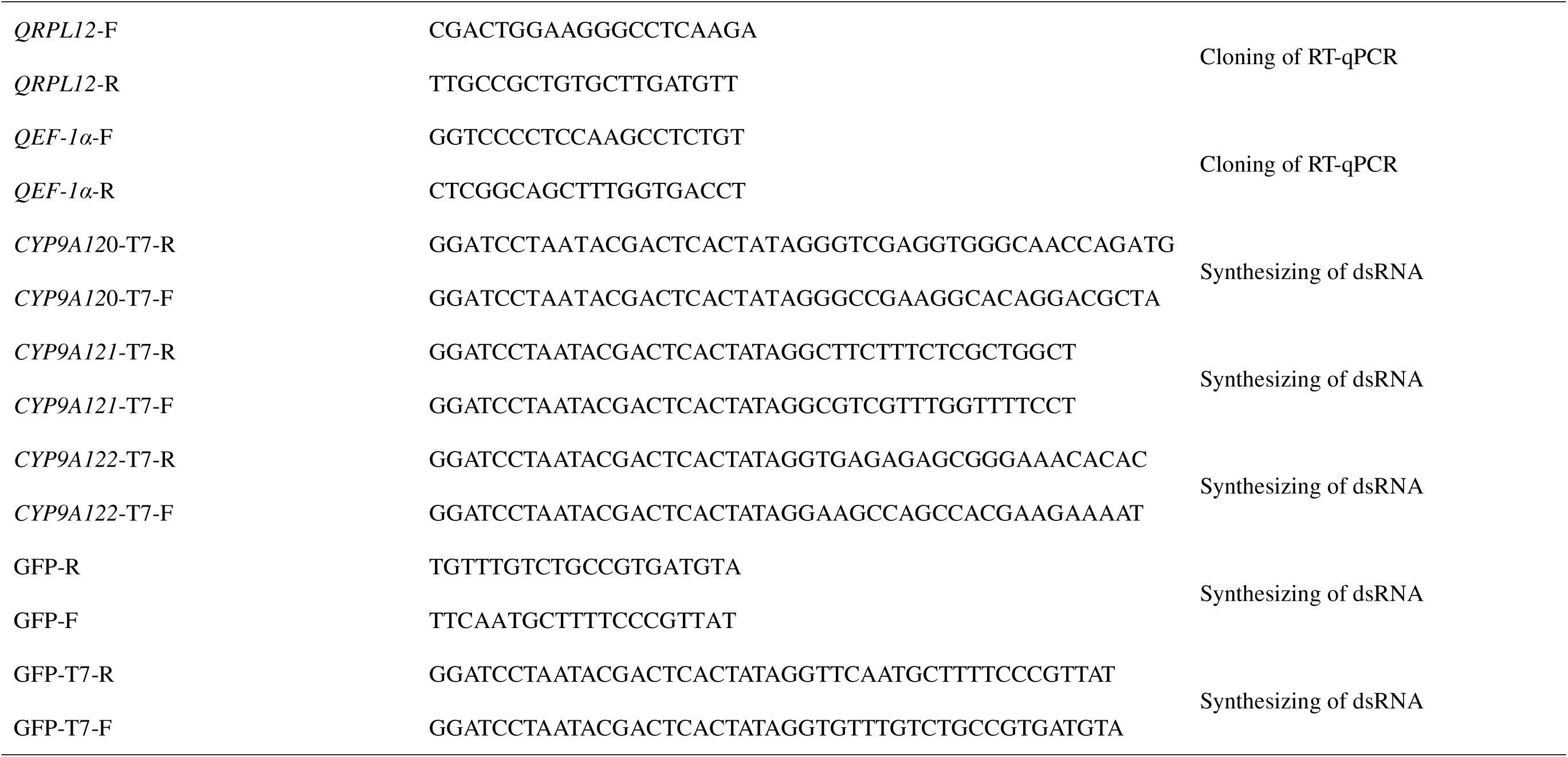
Primers used in this study.

### 2.4 Induction treatment

The dose that would kill 10% (LD_10_) of the larvae by *lambda*-cyhalothrin at 24 h was used for the subsequent insecticide exposure experiment. A 2 μL drop of the insecticide solution was gently dropped on the thoracic dorsum of fourth-instar larvae. Control larvae were treated in the same way with acetone only. After treatment, larvae were held individually in a 2 cm × 5 cm glass tube and fed on an artificial diet. Three replicates, each consisting of 40 surviving larvae were sampled 3, 6, 12, 24, and 48 h post treatment, respectively.

### 2.5 Sampling

A total of 200 eggs, 50 larvae of first- and second-instar, 10 larvae of third-instar, 5 larvae of fourth- and fifth-instar, 5 pupae, and 5 female and male adults, were sampled respectively for each group with three replicates. The head, cuticle, fat body, midgut, and Malpighian tubules were dissected from 30 fourth-instar larvae with three replicates. All of the samples were immediately frozen using liquid nitrogen and then stored at -80 [for subsequent use.

### 2.6 Real-time quantitative reverse transcription PCR (RT-qPCR)

Real-time quantitative reverse transcription PCR (RT-qPCR) was performed following MIQE guidelines^23^ to determine the relative expression levels of *CYP9A* genes. PCR reactions (20 μL) contained 1 μL of 10× diluted cDNA, 10 μL of TB Green Premix Ex Taq 2 (TliRNaseH Plus; Takara, Dalian, China), 0.8 μL of each primer (Table 1). The RT-qPCR was performed on a Bio-Rad CFX96 (Bio-Rad, Hercules, CA, USA) under the thermal cycling conditions comprised 30 s at 95 [, followed by 40 cycles of 95 [for 5 s and 60 [for 30 s. A final melt curve step was included post PCR to determine the specificity of the conduct. The *RPL12* (MT116775) and *EF-1*α (MN037793) were used as reference genes ^18^ for gene expression analysis. Each qPCR contained three biological replicates with three technical repeats for each. The 2 ^-△△CT^ method ^24^ was used to analyze the relative expression of *CYP9A* genes. Data were plotted using GraphPad Prism v5 (GraphPad Software, CA).

### 2.7 dsRNA preparation and silencing of CYP9A by RNAi

To synthesize dsRNA, the T7 RiboMAXTM Express RNAi System (Promega, USA) was used following the manufacturer’s instructions. Forward and reverse primers (Table 1) with a T7 promoter site were designed for PCR amplification. Cycling conditions comprised cycles of 94 [for 30 s, 55 [for 45 s, and 72 [for 1 min, with a final extension step at 72 [for 10 min. PCR amplicons were gel-purified, TA cloned and sequenced as described above. dsRNA was synthesized using T7 RiboMAX^TM^ Express RNAi System (Promega) following the manufacturer′s instructions and was adjusted to a concentration of 3000 ng μL^−1^. *dsGFP* was produced as a control.

The third-instar larvae were used in this study. One μL of dsRNA of either *CYP9A* or *CpGFP* was injected into the thorax of each of the *C. pomonella* larvae. Eighteen larvae were randomly collected at 3, 6, 12, and 24 h after injection. RT-qPCR was performed to determine the reduction in the transcription levels of *CYP9A* genes.

### 2.8 Bioassays and P450 activity after RNAi

The *lambda*-cyhalothrin was dissolved in acetone to produce a stock solution. According to the bioassay result of Wang et al,^5^ 1 μL drop of LD_10_ and LD_50_ dose of *lambda*-cyhalothrin was applied on the thoracic dorsum of each of the larvae from either *CYP9A120* and *CYP9A121* dsRNA-injected or *CpGFP* dsRNA-injected group 6 or 3 h after injection, depending on the persistence of RNAi effectiveness. Fifteen larvae were used for each treatment with three replicates. Mortality was assessed every 12 h for 72 h and results of 48 h after insecticide exposure were compared.

To determine whether silencing of *CYP9A* could effectively affect P450 activity, three replicates of five third-instar larvae of *C. pomonella* at each time interval (3 h, 6 h, 12 h, 24 h, 48 h, and 72 h) after injecting of dsRNA and *dsGFP* were collected. P450 enzyme was prepared as described in Wang et al.^5^ P450 activity was assessed by measuring ECOD activity using 7-ethoxycoumarin as the substrate using the method mentioned in Shi et al.^16^

### 2.9 Heterologous expression of P450s

The ORFs of *CYP9A120* (GenBank No. MF574685), *CYP9A121* (GenBank No. MF574686), *CYP9A122* (GenBank No. MF574692), and NADPH-dependent cytochrome P450 reductase (*CpCPR*, GenBank No. MT159663.1) of *C. pomonella* were synthesized directly after codon optimization (accommodate codon bias of the Sf9 cells from *S. frugiperda*) and inserted to the pFastBac1 vector (Invitrogen). All P450s were heterologously expressed in Sf9 cells with the Bac-to-Bac baculovirus expression system as described.^27^ P450s were co-expressed with CpCPR under a multiplicity of infection (MOI) ratios of 2 and 1, respectively. A non-insertion control was prepared simultaneously under the same co-expression condition. Microsomes of infected cells were purified by differential centrifugation and the total protein content was quantified by the Bradford method.^25^ The recombinant P450 was identified and quantified by a reduced CO-difference spectral assay.^26^

### 2.10 Metabolism of model substrates and *lambda*-cyhalothrin

Four common fluorescent probe substrates were used to identify O-dealkylation and O-debenzylation of recombinant P450s. All tests were conducted at 30 [in 0.1 M potassium phosphate buffer (pH 7.4) with 70 μg microsomes and a final volume of 200 μL. Substrates (10 μM BR, 5 μM BOMR, 25 μM BFC, 25 μM EFC) and microsomes were pre-incubated at 30 [for 5 min, then reactions were started after adding NADPH (10 μL of 10 mM storage buffer) and monitored in real-time mode for 30 min (excitation/emission wavelengths: 530 nm/585 nm for two resorufin substrates and 410 nm/538 nm for two trifluoromethyl coumarin substrates). Reactions were conducted in black, flat-bottom 96-well plates by Spectra Max M5 reader (Molecular Devices, CA, USA). Non-insertion control with an equivalent weight of microsomes was tested simultaneously. The logarithmic phase of each reaction was selected to calculate the final activity and the final metabolic activity was expressed as ΔRFU/min/mg protein. Each experiment has three replicates.

*In vitro* metabolism of *lambda*-cyhalothrin was performed with 30 pmol recombinant P450, NADPH regeneration system (1.3 mM NADP+, 3.3 mM glucose-6-phosphate, 3.3 mM MgCl2 and 0.4 U mL-1 glucose-6-phosphate dehydrogenase) and 50 μΜ *lambda*-cyhalothrin (2 μL freshly dissolved in DMSO) in 200 μL 0.1 M potassium phosphate buffer (pH 7.4). Non-insertion controls and negative control (without NADPH regeneration system) were prepared for each P450 sample and tested simultaneously. Reactions were pre-warmed at 30 for 5 min and started after adding insecticides. Samples were incubated on an orbital shaking incubator at 30 [, 1,200 rpm for 1 h. Reactions were stopped by acetonitrile and purified by centrifugation (15,000 × g) and filtration (0.22 μm). Cleared samples were analyzed immediately by UPLC-MS/MS. All metabolism experiments were repeated twice with three replicates per sample.

### 2.11 UPLC-MS/MS

Samples were separated as described^27^ and briefly as follows: 5 μL samples were separated by Waters Acquity UPLC system (Waters ACQUITY UPLC I-Class) using BEH C18 column (2.1 mm × 50 mm, 1.7 μm particle size. Waters, USA) and eluted with a gradient of mobile phase consisting of acetonitrile (A) and ultrapure water (B), with a constant flow rate of 0.3 mL/min. The gradient elution conditions were as follows: 0 min A: B 40:60, 0.3 min A: B 40:60, 2 min A: B 98:2, 2.5 min A: B 98:2, 2.6 min A: B 100:0, 3 min A: B 100:0, 3.1 min A: B 40:60, 5 min A: B 40:60. For identification of the metabolite, samples separated by UPLC were then analyzed by high-resolution mass spectroscopy (Triple Tof 5600+, AB Sciex, USA) under negative ESI mode. The collision energy for MS/MS was 40 V.

Metabolite formation was regarded as the criterion of enzyme activity for *lambda*-cyhalothrin. The partitioned samples were analyzed using a tandem triple quadrupole mass spectrometer (Waters Xevo TQ-S micro, USA) and run in negative ESI mode. The metabolite was detected under MRM mode (multiple reaction monitoring) with MRM transition: 464 > 205 (quantification) and 464 > 213 (qualification). As the reference compound of the metabolite was not available, the peak area of generated hydroxyl-metabolite was applied for the relative quantification of the catalytic activity of recombinant P450s.

### 2.12 Data analysis

The statistical significance of differences between samples was analyzed with SPSS Statistics 20 (IBM, Chicago, IL, USA). All of the results were presented as mean of triplicates ± standard deviation (SD). Statistical significance was analyzed by one-way analysis of variance (ANOVA) followed by Student’s *t*-test (***, *P* < 0.001).

## 3. RESULTS

### 3.1 Identification CYP9A genes of *C. pomonella*

Apart from the previously reported *CYP9A61*,^7^ three new *CYP9A* genes including *CYP9A120* (GenBank number MF574685), *CYP9A121* (GenBank number: MF574686), and *CYP9A122* (GenBank number MF574692) were identified from *C. pomonella*. The ORF of *CYP9A120*, *CYP9A121,* and *CYP9A122* was 1596, 1602, and 1593 bp, encoding proteins with 531, 533, and 530 amino acids, respectively (Figure S1∼S3). The estimated molecular weight of corresponding deduced proteins of *CYP9A120*, *CYP9A121*, and *CYP9A122* is 61.41, 61.53, and 61.06 kDa, and the theoretical isoelectric point is 7.98, 7.64, and 8.74, respectively (Table 2). These four P450s were all located on chromosome 12 and transcribed from the same strand of DNA (Figure 1A), and shared 54.07%-56.30% identity in amino acid sequence (Table S2). Domain analysis revealed that CYP9A61, CYP9A120, and CYP9A121 all have conserved regions of P450 enzymes including FxxGxxxCxG, PxxFxPxRF, ExxR, and WxxR, while the heme-binding site of CYP9A122 (FxxGXXXCxA) is one residue difference with the others (Figure 1B). Secondary structure analysis revealed that these P450s contain similar amounts of α-helix (ranged from 14 to 16) and β-sheet (ranged from 8 to 10) (Figure S4).

**Figure 1.**
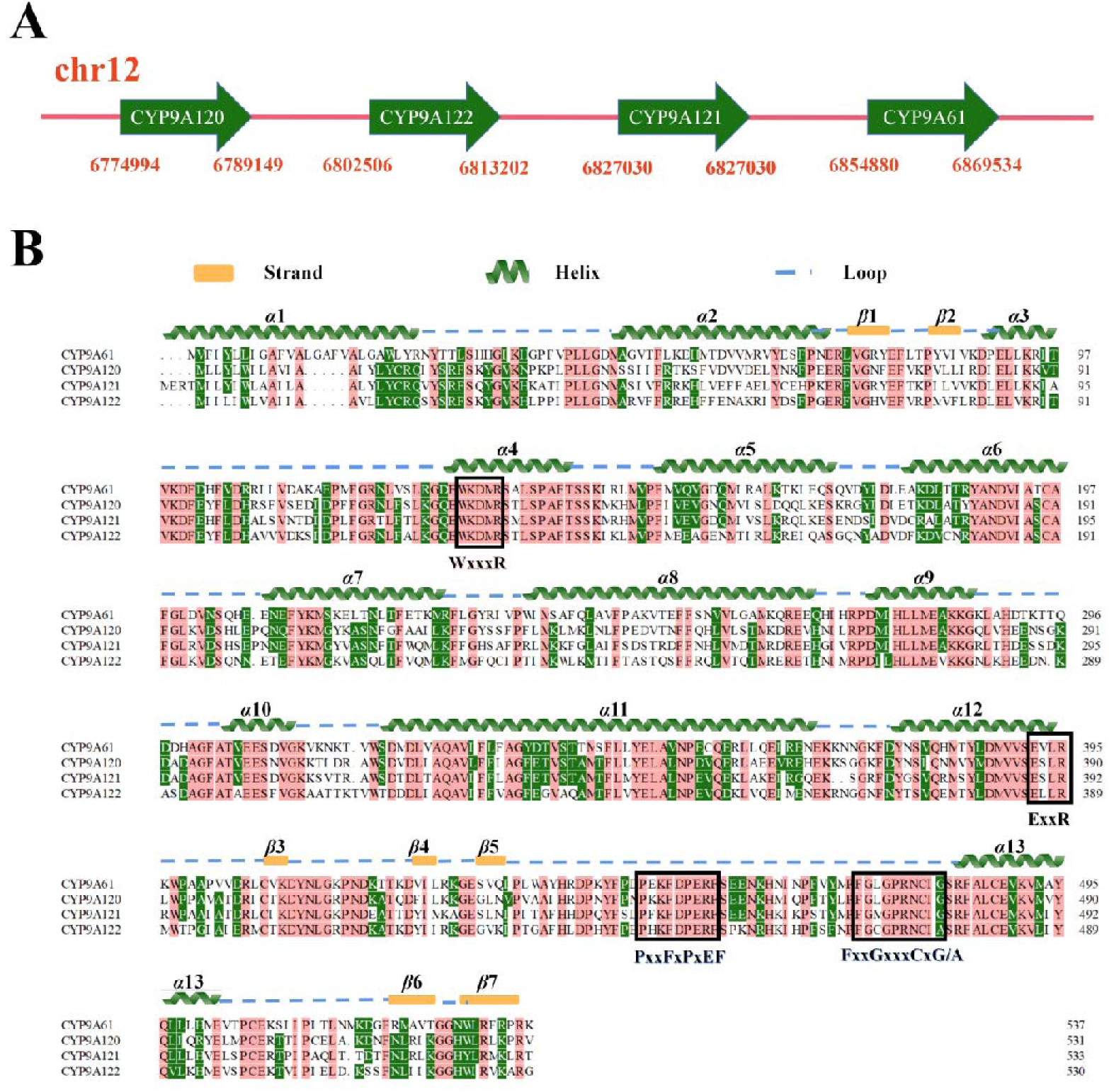
Chromosomal location (A) and alignment of the amino acid sequences of P450s from CYP9A subfamily (B). The conserved regions of P450 enzymes including FxxGxxxCxG/A, PxxFxPxRF, ExxR, and WxxR are boxed. The position of α-helices and β-sheets in the amino acid sequence of CYP9A genes is marked.

**Table 2.** Biological information of CYP9A genes in *C. pomonella*.

| Gene name | ORF (bp/aa) | Mw (KDa) | pI |
| --- | --- | --- | --- |
| *CYP9A61* * | 1617/538 | 62.04 | 7.66 |
| *CYP9A120* | 1596/531 | 61.41 | 7.98 |
| *CYP9A121* | 1602/533 | 61.53 | 7.64 |
| *CYP9A122* | 1593/530 | 61.06 | 8.74 |

\*Note: Biological information of *CYP9A61* gene are detailed in Yang et al.<sup>17</sup>

The neighbor-joining tree revealed that the *CYP9A* subfamily from *C. pomonella* contains three clades (Figure 2). *CYP9A120* and *CYP9A121* clustered together within a branch with strong bootstrap support *CYP9A122* but separated from *CYP9A122* and *CYP9A61*, which might be the result of an ancient duplication with separating the *CYP9A* lineage from the other *CYP9A* genes. Although located on the same chromosome, *CYP9A61* clusters near *CYP9A3* from *H. armigera*, suggesting that *CYP9A61* is a probable ortholog of *H. armigera CYP9A3* (Figure 2).

**Figure 2.**
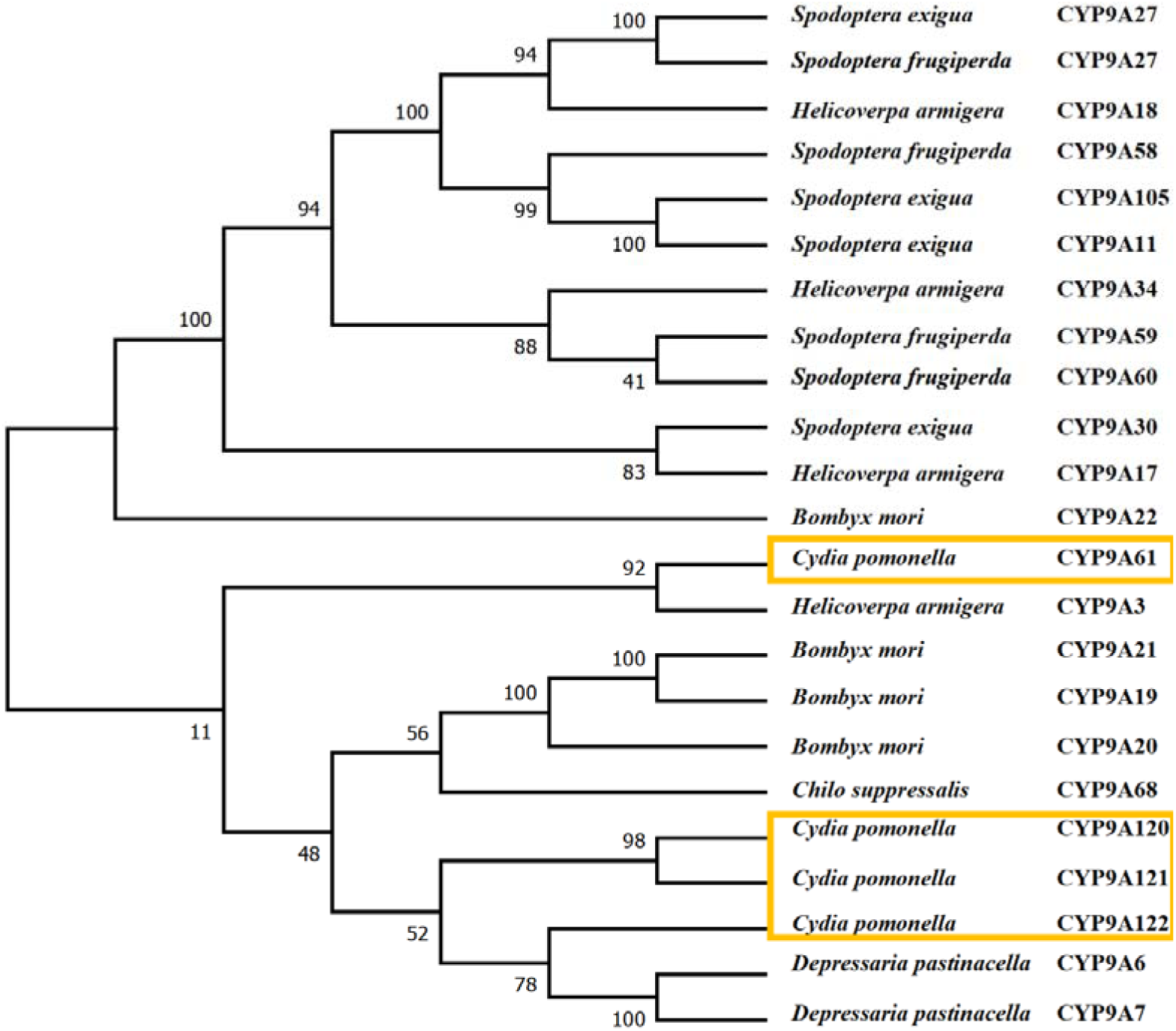
Phylogenetic analysis of *CYP9A* genes from *C. pomonella*, *H. armigera*, *B. mori*, *S. frugiperda*, *S. exigua*, *Depressaria Pastinacella* and *C. suppressais*. The *CYP9A* genes from *C. pomonella* are boxed. Information for P450 used for phylogenetic analysis are shown in Table S4.

### 3.2 Development stage and tissue-specific expression profiles of P450 genes

Spatio-temporal expression pattern showed that CYP9A genes were expressed in all developmental stages and tissues. Developmental stage expression profiling of *CYP9A120* (*df*=9, *F*=72.59, *p*<0.001), *CYP9A121* (*df*=9, *F*=21.31, *p*<0.001), and *CYP9A122* mRNA transcripts revealed higher expression levels during the larval stage (*df*=9, *F*=27.53, *p*<0.001). *CYP9A120* was mostly expressed in the fourth instar, while *CYP9A121* and *CYP9A122* were most abundant in the first- and fifth-instar of larvae, respectively (Figure 3A). Tissue-specific expression analysis revealed that these three *CYP9A* genes were most highly expressed in the midgut (*CYP9A120*: *df*=4, *F*=157.24, *p*<0.001; *CYP9A121*: *df*=4, *F*=66.77, *p*<0.001; *CYP9A122*: *df*=4, *F*=73.88, *p*<0.001), which indicates that these gut-specific genes may play a key role in detoxification processes (Figure 3B).

**Figure 3.**
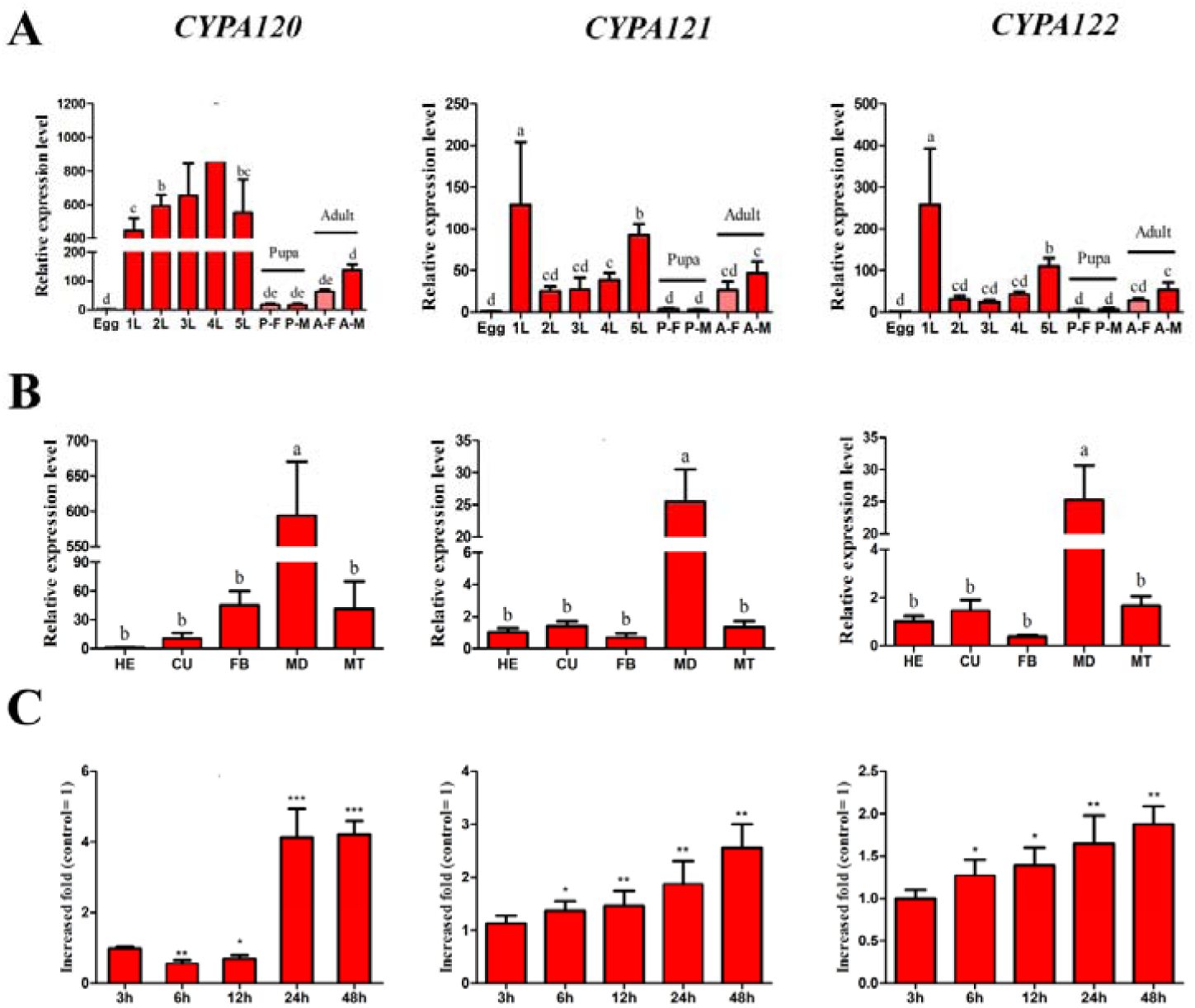
The expression level of *CYP9A* genes in *C. pomonella*. (A) developmental stage expression profiles. P-F, pupa female; P-M, pupa male; A-F, adult female; A-M, adult male. (B) Tissue-specific expression profiles. HE, head; CU, cuticle; FB, fat body; MD, midgut; MT, Malpighian. (C) LD_10_*lambda*-cyhalothrin induced expression profiles. The data was used the mean ± SD and the error bars stand for the standard errors derived from the three replicates. The letters on the error bars show the significant difference among the different development stages and tissues analyzed by the one-way analysis of variance (ANOVA) with Duncan’s test (*P* < 0.05). The asterisks on the error bars show the significant difference among the different treatment times analyzed by the one-way analysis of variance (ANOVA) with the Student’s *t*-test (***, *P* < 0.001).

### 3.3 Expression of CYP9A genes exposed to *lambda*-cyhalothrin

Exposure of fourth-instar larvae to LD_10_ of *lambda*-cyhalothrin significantly increased the transcription levels of *CYP9A121* and *CYP9A122* over time (2.53- and 1.87-fold), whereas the expression level of *CYP9A120* was decreased at 6 and 12 h after exposure, and was then increased by 4.21-fold at 24 h post exposure (Figure 3C).

### 3.4 Expression profiles of CYP9A genes in field-evolved *lambda*-cyhalothrin-resistant population

Transcript levels of *CYP9A120*, *CYP9A121,* and *CYP9A122* genes were 41.56-, 4.55-, and 2.73-fold higher, respectively, in the ZW_R population in comparison to the laboratory susceptible strain SS strain (Figure 4).

**Figure 4.**
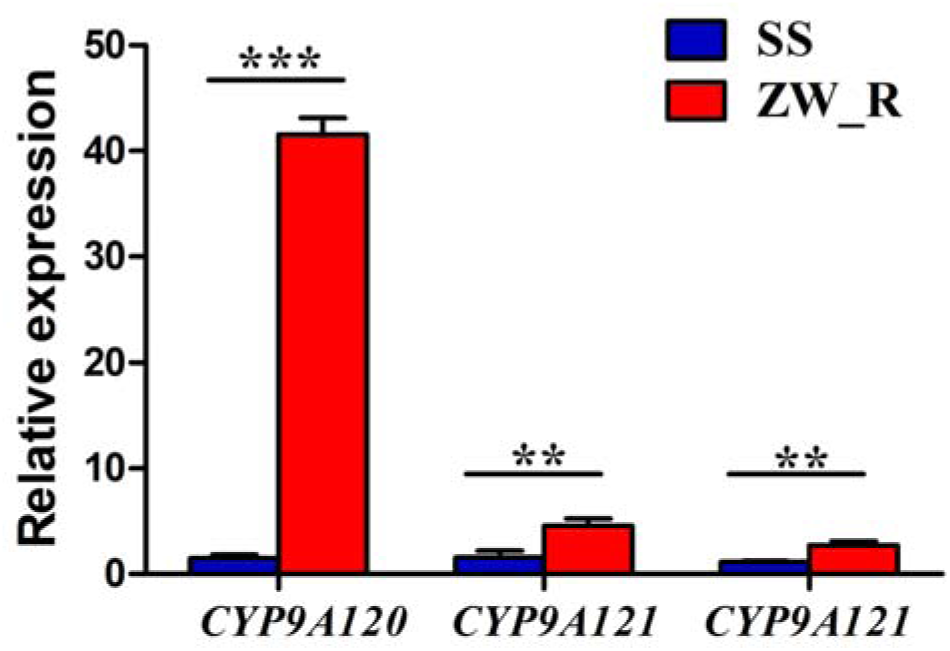
Expression profiles of *CYP9A* genes in the field *lambda*-cyhalothrin resistant population of *C. pomonella*. The results are shown as the mean ± SD. The error bars represent the standard errors calculated from three replicates. Asterisks on the error bar indicate significant differences analyzed by the one-way analysis of variance (ANOVA) with the Student’s *t*-test (***, *P* < 0.001).

### 3.5 RNAi efficacy and susceptibity to *lambda*-cyhalothrin

The expression level of *CYP9A120*, *CYP9A121,* and *CYP9A122* genes decreased dramatically at 3 h after injection of *dsCYP9A120*, *dsCYP9A121*, and *dsCYP9A12* compared with *dsGFP* (Figure 5A). The RNAi efficacy was 49.76% ± 14.49%, 59.67% ± 9.87% and 52.13% ± 10.62% for *dsCYP9A120* (Figure 5A), and 68.39% ± 15%, 53.75% ± 12% and 20.09% ± 14.2% for *dsCYP9A121* (Figure 5B) at 3 h, 6 h and 12 h after injection of dsRNA, respectively. The effect has only lasted for 3 h in larvae injected with *dsCYP9A122* (52.15% ± 8.77%) compared with *dsGFP* (Figure 5C). The ECOD activity was suppressed after silencing *CYP9A120* and *CYP9A121* in comparison to dsGFP, while *CYP9A122* did not significantly affect the activity of P450 (Figure S5).

**Figure 5.**
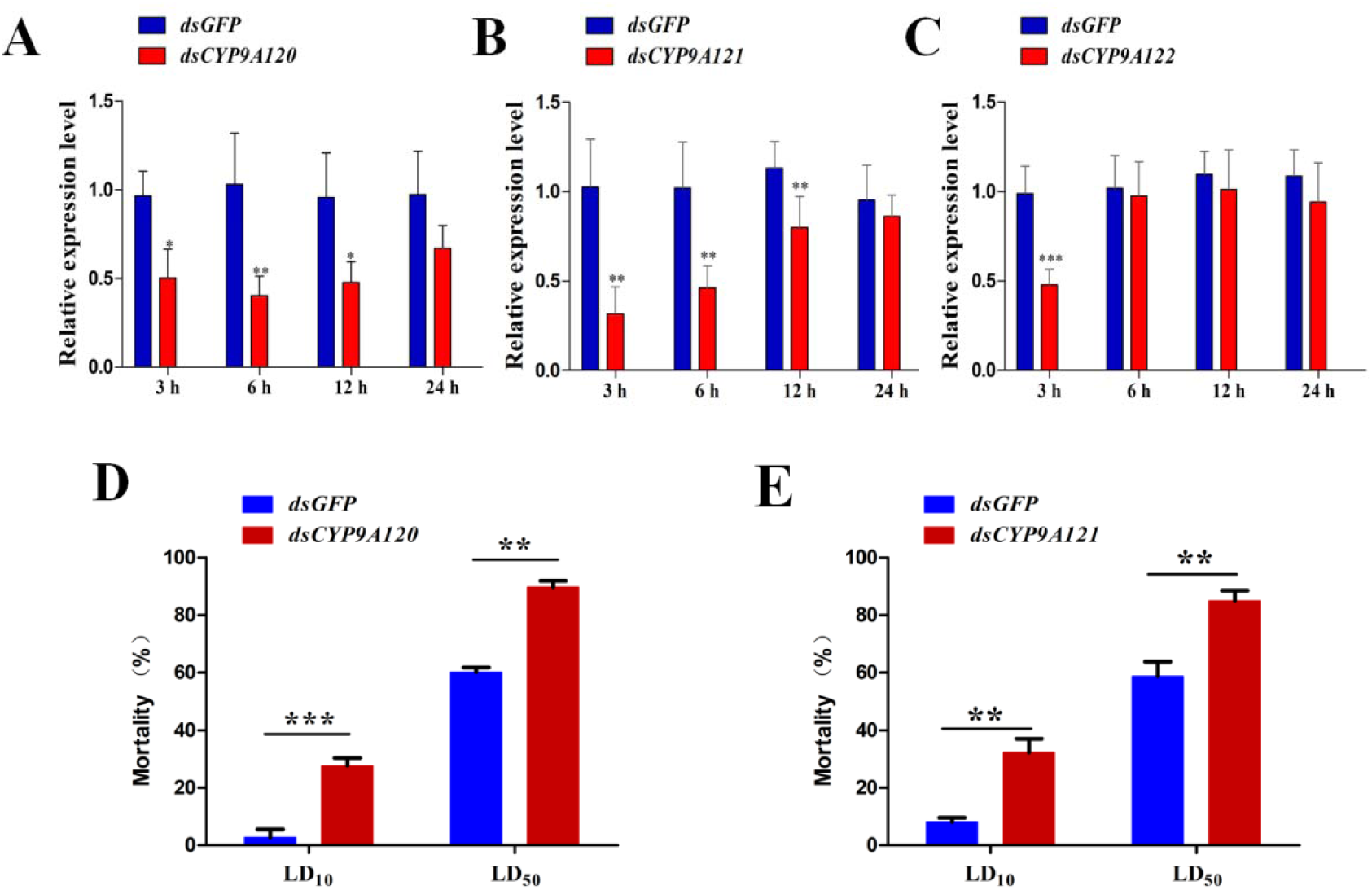
Functional analysis of *CYP9A* genes in *C. pomonella* by RNAi. (A) Relative expression levels of *CYP9A120* (A), *CYP9A121* (B), *CYP9A122* (C) in fourth-instar larvae of *C. pomonella* injected with *dsGFP* or *dsCYP9As*. (D) The mortality of fourth-instar larvae of *C. pomonella* injected with *dsCYP9A120* (D) and *dsCYP9A121* (E) was assessed every 12 h after the insecticide treatment in comparison to those injected with *dsGFP*. The results shown are expressed as means ± SE of three biological replicates. Asterisks on the error bar indicate significant differences analyzed by the one-way analysis of variance (ANOVA) with the Student’s *t*-test (\**P* < 0.05, ** *P* < 0.01, *** *P* < 0.001).

To examine whether the knockdown of certain gene induces off-target effects, transcript levels of other *CYP9A* genes in larvae after dsRNA injection were analyzed. The result showed that RNAi induced off-target effects were not observed, and the expression levels of CYP9A subfamily genes were not affected by silencing *CYP9A120* and *CYP9A121* (Figure S6).

Knockdown of *CYP9A120* and *CYP9A121* in the larvae of *C. pomonella* significantly increased their susceptibility to *lambda*-cyhalothrin compared with the control (*P* < 0.05). Bioassays showed that 27.77% and 87.87% of *CYP9A120*-knockdown larvae died after exposure to LD_10_ and LD_50_ of *lambda*-cyhalothrin, respectively (Figure 5D). Similarly, the sensitivity of *CYP9A121* dsRNA-injected larvae to LD_10_ and LD_50_ of *lambda*-cyhalothrin was enhanced, with mortality of 32.28% and 84.97%, respectively, which was significantly higher than the control groups (Figure 5E).

### 3.6 Functional expression of CYP9As in Sf9 cells

The *CYP9A120*, *CYP9A121,* and *CYP9A122* were expressed in Sf9 cells using the baculovirus system. Reduced CO-difference spectra of all recombinant P450 proteins in purified microsomes showed a maximum peak near 450 nm (Figure 6), indicating successful expression of functional enzymes within Sf9 cells. Among four tested fluorescent substrates, three CYP9As showed activities to both BFC (Figure 7C) and BOMR (Figure 7D), again indicating that the recombinant enzymes had correct folding and were active.

**Figure 6.**
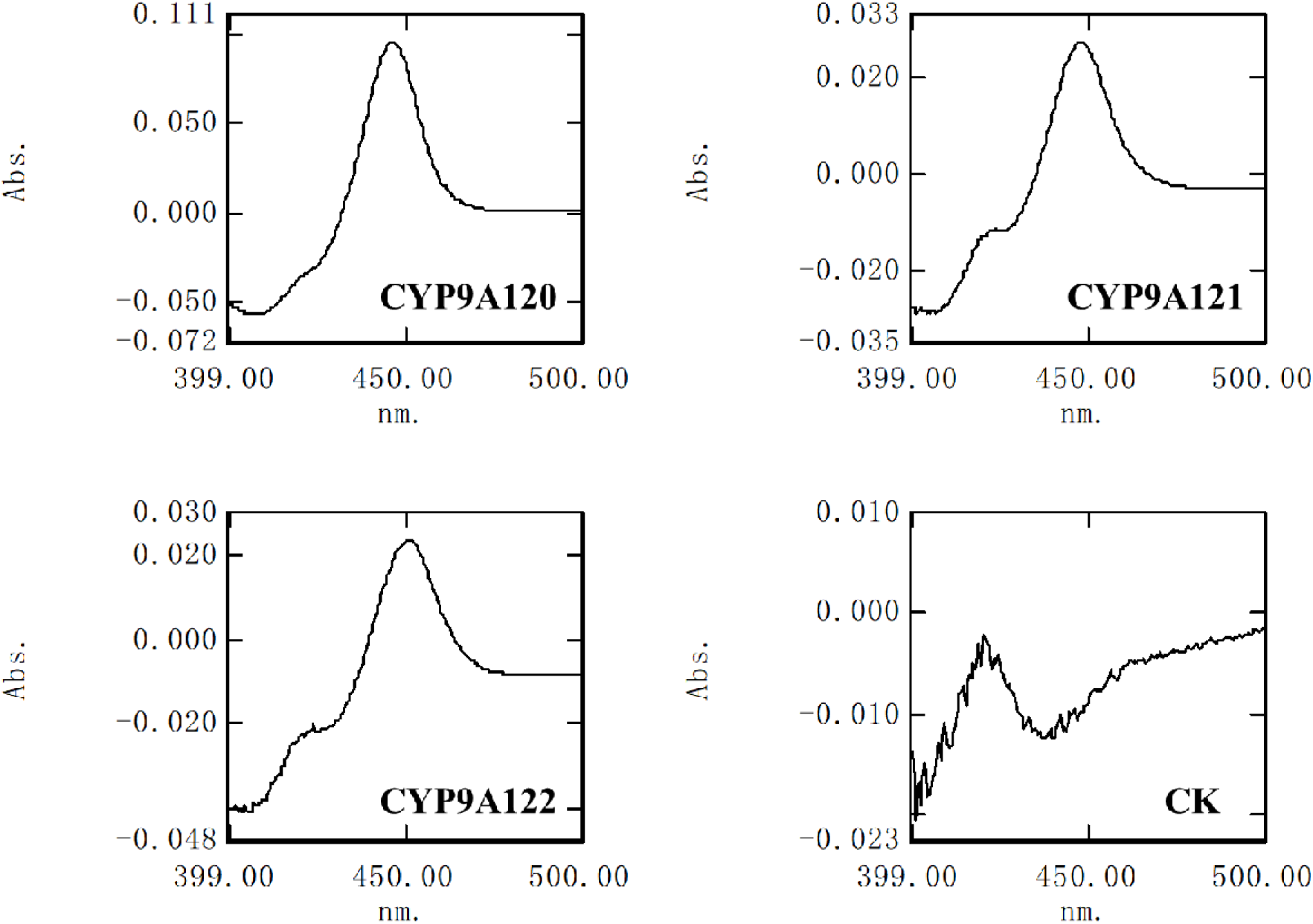
Reduced CO-difference spectra of three recombinant CYP9A proteins. Sf9 cells expressing individual P450s were lysed and subjected to co-difference spectral tests. Sf9 cells infected with the non-insertion virus were used as negative control (CK).

**Figure 7.**
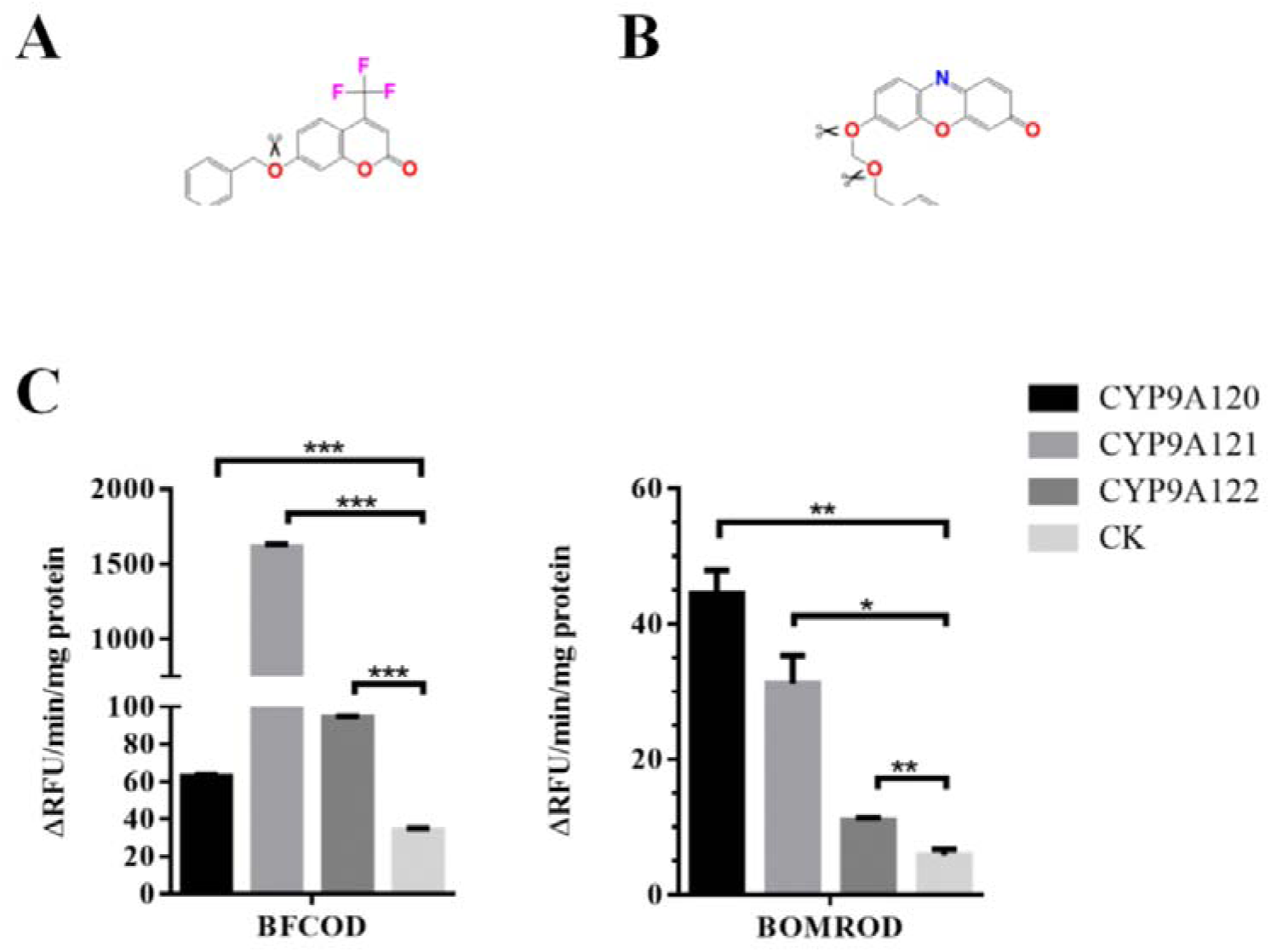
Metabolism of two artificial fluorogenic substrates by three recombinant CYP9A proteins. (A) Structure of BFC. (B) Structure of BOMR. (C) Metabolism of BFC and BOMR by three recombinant CYP9A proteins. Data are mean values ± SEM (n=4). Asterisks on the error bar indicate significant differences analyzed by the one-way analysis of variance (ANOVA) with the Student’s *t*-test (\**P* < 0.05, ** *P* < 0.01, *** *P* < 0.001).

### 3.7 Metabolism of *lambda*-cyhalothrin

Two distinct metabolites (M+16) with a retention time of 2.24 ± 0.02 min and 2.31 ± 0.01 min, respectively, were detected in recombinant CYP9As samples (Figure 8A). Both metabolites showed a similar quasi-molecular ion and specific isotope peak ratio of 3:1 (n: n+2) in the full scan spectrum. The highest isotope peak of the two metabolites was 464.0877 (2.26 min, error = -1.1 ppm) and 464.0882 (2.32 min, error= 0 ppm [∼0.01 mDa]), respectively. Moreover, typical fragments of chrysanthemic acid group (m/z 205/241) and oxidized henoxybenzyl alcohol group (m/z 185/213) were found by MS/MS (Figure 8B). In general, these two metabolites were hydroxy-metabolites of *lambda*-cyhalothrin. Furthermore, a specific fragment of 2′-OH-metabolite (m/z = 222) was found in the MS/MS spectrum of hydroxyl-metabolite at 2.32 min. According to the published information of hydroxyl-metabolites of fenvalerate,^16^ the former metabolite (2.26 min) might be 4′-OH-*lambda*-cyhalothrin and the latter (2.32 min) might be 2′-OH-*lambda*-cyhalothrin. For highly sensitive detection of metabolites, the most abundant isotopic ion 464 and fragment 205 were used as parent ion and daughter ion for MRM detection.

**Figure 8.**
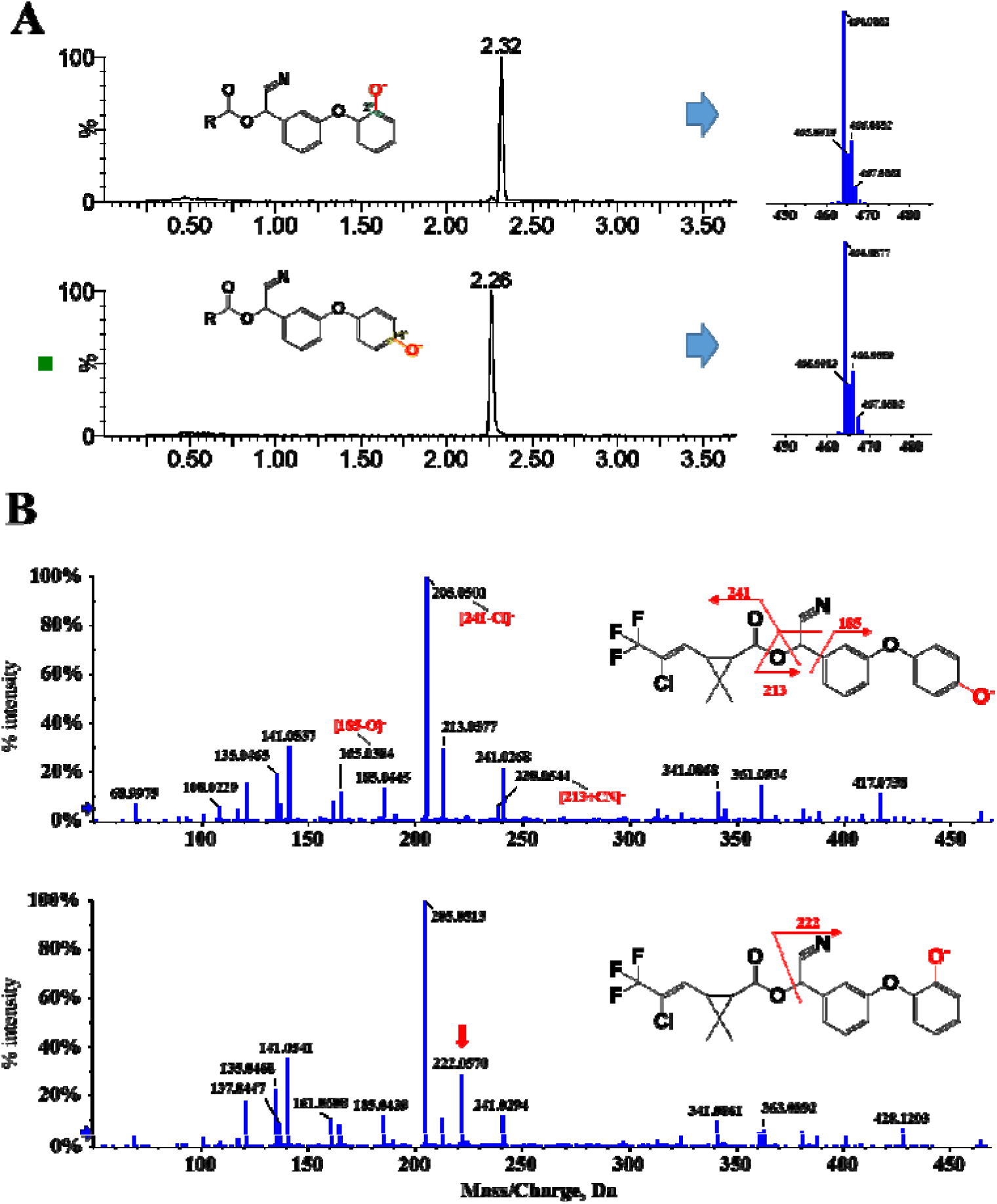
Identification of the hydroxy metabolites of *lambda*-cyhalothrin. (A) Extracted ion currents and MS spectrums of two hydroxy-metabolite. (B) ESI-TOF high-resolution MS/MS spectrum of two metabolites. The fragmentation patterns of the key daughter ions are shown at the top right of each spectrum.

*In-vitro* metabolism showed that *lambda*-cyhalothrin can be metabolized into hydroxy-metabolites by recombinant CYP9As. No hydroxy-metabolites were found in samples without NADPH (-NADPH) (Figure 9A). Meanwhile, samples with recombinant CYP9As generated significantly more metabolites than non-insertion control (CK) (Figure 9C). The 4’-hydroxylated metabolites have been regarded as the main metabolite of pyrethroids metabolized by P450 in *vitro*,^28^ and was detected as the only metabolite of *lambda*-cyhalothrin from CYP9A121. However, the 2’-OH-metabolites-*lambda*-cyhalothrin, another but the only metabolite of *lambda*-cyhalothrin was formed by CYP9A122. For CYP9A120, both 2′- and 4′-metabolites were generated (Figure 9A & 9C).

**Figure 9.**
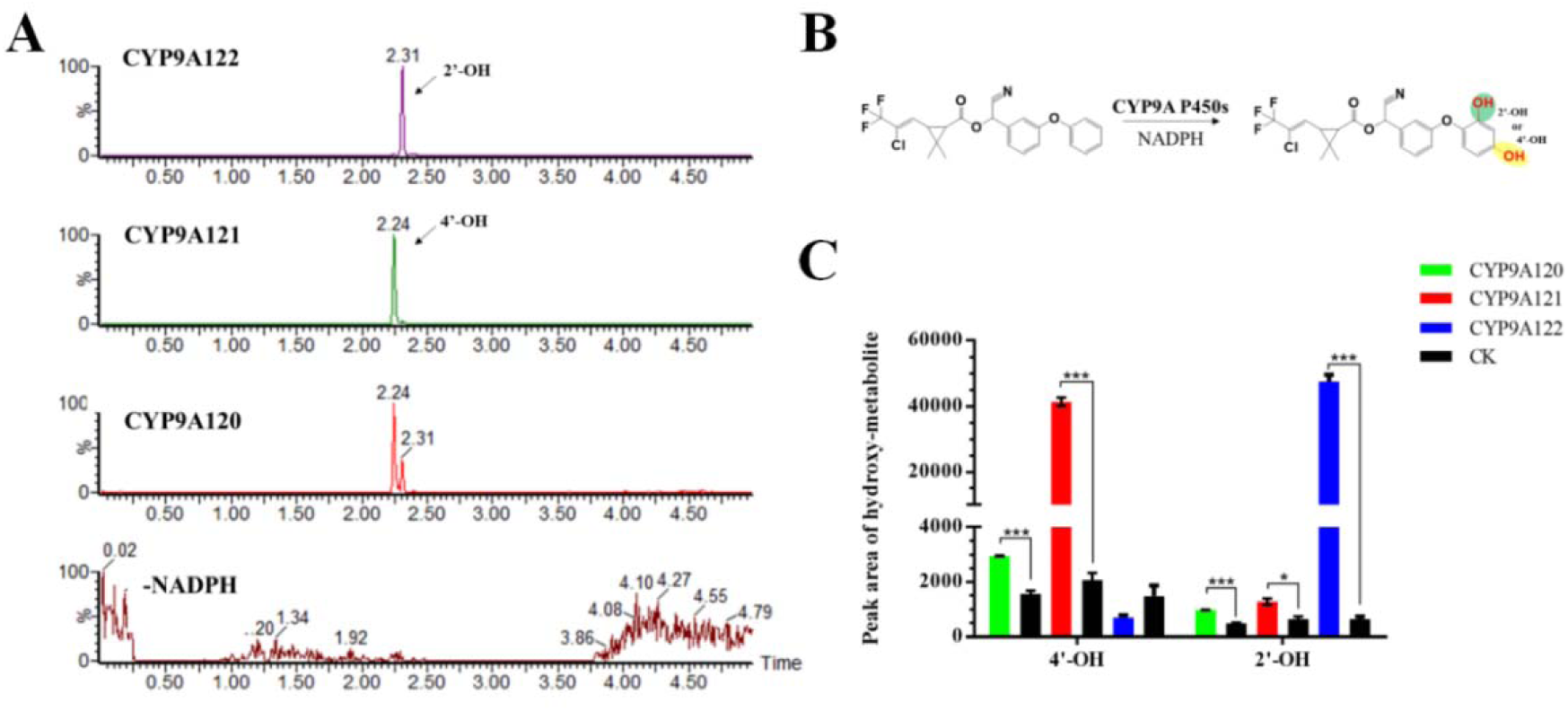
Metabolism of *lambda*-cyhalothrin by three recombinant CYP9A proteins. (A) MRM signals of samples with CYP9A120, CYP9A121, CYP9A122, and without NADPH (-NADPH). (B) The sketch map of recombinant CYP9A metabolized *lambda*-cyhalothrin. (C) Relative quantification of metabolic competencies of three CYP9A P450s. Data are mean values ± SEM (n=3-7). Asterisks on the error bar indicate significant differences analyzed by the one-way analysis of variance (ANOVA) with the Student’s *t*-test (\**P* < 0.05, *** *P* < 0.001).

## 4. DISCUSSION

The use of synthetic insecticides for pest control protects crops but imposes strong selection pressures to pests that can lead to the development of resistance,^29, 30^ and may also result in pest outbreaks.^31, 32^ Apart from the alterations in target sites that prevent insecticides from binding to them,^33^ the ability of herbivorous insects to detoxify insecticides is another common resistance mechanism and is pivotal for their adaptive evolution^34^ although often linked with fitness costs.^35–37^ P450s especially members from the *CYP9* family usually contributed to pyrethroids resistance in Lepidoptera pests,^16, 38, 39^ but the molecular mechanism of metabolic resistance caused by P450 genes in *CYP9* family is still unclear in *C. pomonella*. In this study, we showed that overexpression of *CYP9A* subfamily genes likely contributed to *lambda*-cyhalothrin resistance in *C. pomonella*.

Four members of the *CYP9A* subfamily, including *CYP9A61*, *CYP9A120*, *CYP9A121*, and *CYP9A122* were identified and located on *C. pomonella* chromosome 12. No premature stop codons were found in either sequence and contain conserved motifs of P450, suggesting that these genes are not a pseudogene and are likely to be functional. The protein sequence of these newly identified P450 genes (*CYP9A120*, *CYP9A121*, and *CYP9A122*) share 55.564%, 56.30%, and 54.07% identity with *CYP9A61* (Table S2), known to be a potential metabolizer of *lambda*-cyhalothrin and chlorpyrifos-ethyl in this species.^40^ The great diversity of P450 genes in arthropods is well documented and the number of *CYP* genes in each species is highly variable.^41^ In this study, we found that the number of *CYP9A* genes of *C. pomonella* is the same as that of *B. mori a*nd *C. suppressalis*, but less than that of *Spodoptera* species (*S. exigua*, *S. frugiperda*, *S. litura*, *P. xylostella*, and *H. armigera*) (Table S3). This is mainly because *S. litura* and *S. exigua* are the typical polyphagous pests, and *S. frugiperda* can harm almost all gramineous crops. They have evolved P450s equipped with more *CYP9A* genes that can detoxify xenobiotics, including secondary plant substances and insecticides they encounter in their environment.^42^ In addition, *CYP9A120*, *CYP9A121,* and *CYP9A122* probably have a similar function in metabolizing insecticides because they clustered in the same clade with *CYP9A68* from *C. suppressalis*^43^ and *CYP9A19* and *CYP9A22* from *B. mori*,^44^ which have been documented with roles in the development of resistance to triazophos and phoxim, respectively. These results indicate that the *CYP9A* genes may have a potential role in the detoxification of and tolerance or resistance to xenobiotics in *C. pomonella*.

Insect CYP genes are expressed in almost all tissues and throughout most developmental stages. Therefore, tissue-specific and developmental expression profiles of CYP genes may provide insights into their biological and physiological functions.^45^ Previous work on the *CYP9A61* in *C. pomonella* revealed a high level of expression in the fourth- and fifth-instar of larval, which is the main feeding period for *C. pomonella*.^7^ Although *CYP9A120*, *CYP9A121,* and *CYP9A122* genes were ubiquitously expressed throughout all developmental stages of *C. pomonella*, their expression profiles are not identical. The relative expression level of *CYP9A120* in the whole larval stage is higher than in pupal and adult stages, while *CYP9A121* and *CYP9A122* were mainly expressed in the first-instar larvae (neonate larvae), which is the main target of insecticides on *C. pomonella*. Moreover, transcript levels of *CYP9A121* and *CYP9A122* in fifth-instar larvae were significantly higher than in second-, third-, and fourth-instar larvae. Similar developmental expression patterns have been reported in *CYP* genes from *C. suppressalis*,^46^ and *Culex quinquefasciatus*.^47^ As the main stage of feeding and target of insecticides, the high expression level of P450 genes in the larval stage of *C. pomonella* indicates that these genes may be involved in the metabolism of plant allelochemicals and insecticides.

The midgut is the organ for digestion of food and detoxification of xenobiotics in insects, and *CYP* genes expressed in the midgut likely act to detoxification of insecticides.^7^ For example, *CYP9A17* is predominantly expressed in the larval midgut of *H. armigera*, and is responsible for the detoxification of deltamethrin, gossypol, phenobarbital, and capsicum.^48^ In this study, *CYP9A120*, *CYP9A121,* and *CYP9A122* were highly expressed in the midgut of *C. pomonella*. Unlike *CYP9A120*, *CYP9A121,* and *CYP9A122*, *CYP9A61* was highly expressed in the fat body, which is a tissue with a known role in xenobiotic detoxification.^7^ These results imply that *CYP9A* genes are likely responsible for the detoxification of xenobiotics including insecticides and plant allelochemicals in *C. pomonella*.

P450s play a key role in the detoxification of xenobiotic and endogenous compounds, which cause a high-level resistance and adaptation to synthetic insecticides in insects.^15, 28, 49^ Induction of arthropods’ P450 genes in response to insecticides they encounter in their environment is one of the adaptive strategies to defend themselves,^50, 51^ but provides the only evidence of potential P450 involvement in insecticide detoxification. *Lambda*-cyhalothrin is a pyrethroids insecticide with a high insecticidal activity that is widely used for the control of *C. pomonella*.^5^ Our previous work revealed that *CYP9A61* is involved in the detoxification of *lambda*-cyhalothrin. To determine the role of other *CYP9A* genes responsible for the detoxification of *lambda*-cyhalothrin, we investigated the expression levels of *CYP9A120*, *CYP9A121,* and *CYP9A122* in response to a sublethal dose of *lambda*-cyhalothrin. The expression levels of these *CYP9A* genes were induced by LD_10_ of *lambda*-cyhalothrin. Up-regulation of P450 genes by insecticides has been reported in several insect species. For example, *CYP9A59* was effectively induced by methoxyfenozide in the midgut of *S. frugiperda*.^52^ Deltamethrin can significantly increase the expression level of *CYP9A40* in the midgut and fat body of *S. litura*.^53^ Induction of *CYP9A* genes is also observed and is likely responsible for the detoxification of pyrethroids in *Locusta migratoria*.^54^ These results suggest that *CYP9A* genes may play a role in *lambda*-cyhalothrin detoxification in *C. pomonella*. The first *CYP9A* was found in a thiodicarb-resistant population of *Heliothis virescens* and named *CYP9A1*.^55^ Thereafter, several studies have documented that over-expression of *CYP9As* is associated with insecticide resistance.^56^ To further investigate the roles of *CYP9A* genes in *lambda*-cyhalothrin resistance, the relative expression of *CYP9A120*, *CYP9A121*, and *CYP9A120* was compared between resistant and susceptible populations. All *CYP9A* genes were overexpressed in ZW_R, a field population that has evolved a moderate level of resistance to *lambda*-cyhalothrin.^18^ Of these, the relative expression of *CYP9A120* in the resistant population was much higher than the others, suggesting it might play a more important role in resistance. Similar results that overexpression of P450 genes in insecticide-resistant populations have also been reported in several insect species, including *C. suppressalis* ^46^ and *H. armigera*.^57^ These results imply that overexpression of *CYP9A* genes was associated with *lambda*-cyhalothrin resistance in *C. pomonella*.

Numerous investigations have shown that several P450 genes exhibited a distinct molecular response at the transcriptional level when exposed to insecticides, however, this is insufficient systematic functional analyses such as metabolism *in vitro*, RNAi, and CRISPR-Cas9 should be involved.^58^ To further investigate the role of *CYP9A* genes in *lambda*-cyhalothrin resistance, the loss-of-function experiment was performed by RNAi. We employed an injection-based RNAi method that had been used in *S. litura*^59^ to knock down *CYP9A* genes. When the expression of *CYP9A120* and *CYP9A121* was suppressed by injection of dsRNA, the mortality of larvae treated with the LD_10_ and LD_50_ of *lambda*-cyhalothrin increased significantly. Similarly, RNAi mediated silencing of *CYP9A105* caused a significant increase in susceptibility of *S. exigua* larvae to α-cypermethrin, deltamethrin, and fenvalerate,^60^ and suppression of *CYP9A3* and *CYP9AQ1* expression lead to increased mortality of *L. migratoria* to deltamethrin and fluvalinate.^54^ Simultaneously, after the silencing of *CYP9A120* and *CYP9A121* by RNAi, the total activity of P450 was significantly reduced, suggesting these genes encode the functional enzyme. In comparison to *CYP9A120* and *CYP9A121*, the RNAi efficiency of *CYP9A122* was much lower, probably due to a relative lower expression level of this gene (Figure S7). These results are consistent with the opinion that Lepidoptera has much lower RNAi efficiencies^61^ and the sensitivity of an insect to dsRNA varies drastically due to many factors.^62^ These results indicate that *CYP9A120* and *CYP9A121* are responsible for *lambda*-cyhalothrin resistance in *C. pomonella*.

To further confirm the role of these P450s are responsible for the degradation of and resistance to *lambda*-cyhalothrin, we determined the catalytic activity of the P450s by measuring the production of hydroxy-metabolites of *lambda*-cyhalothrin. The result indicated that CYP9A120, CYP9A121, and CYP9A122 can metabolize *lambda*-cyhalothrin. Considering the previous research in which CYP9A was responsible for the detoxification of and resistance to another pyrethroid insecticide, esfenvalerate in *H. armigera*,^16, 63^ we concluded that CYP9A subfamily genes likely play an important role in resistance to pyrethroid insecticides in insects. In this study, we found regioselectivity of metabolism of *lambda*-cyhalothrin for different members of the CYP9A subfamily. CYP9A122 preferred to catalyze ortho-position of the outside benzene ring and only generated 2′-OH-metabolite. In contrast, the metabolite with hydroxylation at para-position (4′-OH-metabolite) was much more than 2′-OH-metabolite for CYP9A121. Nevertheless, CYP9A120 maintained low but relatively balanced catalytic competence to generate both 2′- and 4′-metabolites. Our results combined with previous data^16^ show that CYP9As can not only convert pyrethroids to 4’-hydroxylated but also 2’-hydroxylated metabolites. Similarly, ten P450s from CYP6B and CYP9A subfamilies can efficiently metabolize esfenvalerate into 4’-hydroxy-metabolite, but 2’-hydroxylated was also produced by CYP9A12.^16^ These results demonstrated that *CYP9A* genes in *C. pomonella* have a considerable degree of functional redundancy in terms of *lambda*-cyhalothrin metabolism, which might be a result of evolutionary divergence and functions of P450. Therefore, the serious resistance to insecticides globally in *C. pomonella* is more easily understood, which is probably associated with the subfunctionalization of *CYP9A* genes. The functional redundancy of *CYP9A* genes also challenges the sustainable management of this invasive species using chemical insecticides, as the application of insecticides may lead to growing selection pressure for *C. pomonella* and increase the expression of *CYP9A* genes that contribute to resistance. It is well known that changes leading to P450-mediated insecticide resistance may result from quantitative changes in the expression of P450 that increased the number of the P450s able to metabolize insecticide, as well as qualitative changes in P450 coding sequences.^15^ Several studies have demonstrated that quantitative changes by duplication or amplification of P450 are the main driving force for the evolution of insecticide resistance.^64–67^ Recently, a study in *S. frugiperda* suggested that resistance to deltamethrin is mainly due to the copy number variation of *CYP9A* genes.^68^ Therefore, more work including quantitative and qualitative changes of P450 is needed to fully understand the molecular mechanisms of P450-mediated resistance to insecticide in *C. pomonella*. Taken together, our results provide evidence that the overexpression of *CYP9A* genes is partly responsible for resistance to *lambda*-cyhalothrin in *C. pomonella*. These findings improve our understanding of the evolution of P450 genes, and not only providing new insights into the mechanisms of P450-mediated resistance to insecticides and resistance management of *C. pomonella*, but also for elucidating the mechanisms of global spread of this invasive species.

## ACKNOWLEDGEMENT

This work was supported by the National Key R&D Program of China (2021YFD1400200), and the National Natural Science Foundation of China (31972299).

## Supporting information

**Table S1.** Information of amino acid sequences of CYP9A used as queries for searches of TBLASTN.

| Insect species | Gene name | Accession number |
| --- | --- | --- |
| *Helicoverpa armigera* | *CYP9G5* | KM016757.1 |
|  | *CYP9A12* | EU541248.2 |
|  | *CYP9A14* | KR095603.1 |
|  | *CYP9A17* | AY753201.1 |
| *Spodoptera exigua* | *CYP9A9* | AB381883.1 |
| *Bombyx mori* | *CYP9A19* | EF488994 |
|  | *CYP9A20* | EF421989 |
|  | *CYP9A21* | EF415298.1 |
| *Mamestra brassicae* | *CYP9A90* | KR676343.1 |
| *Cydia pomonella* | *CYP9A61* | KC832920 |

**Table S2.**
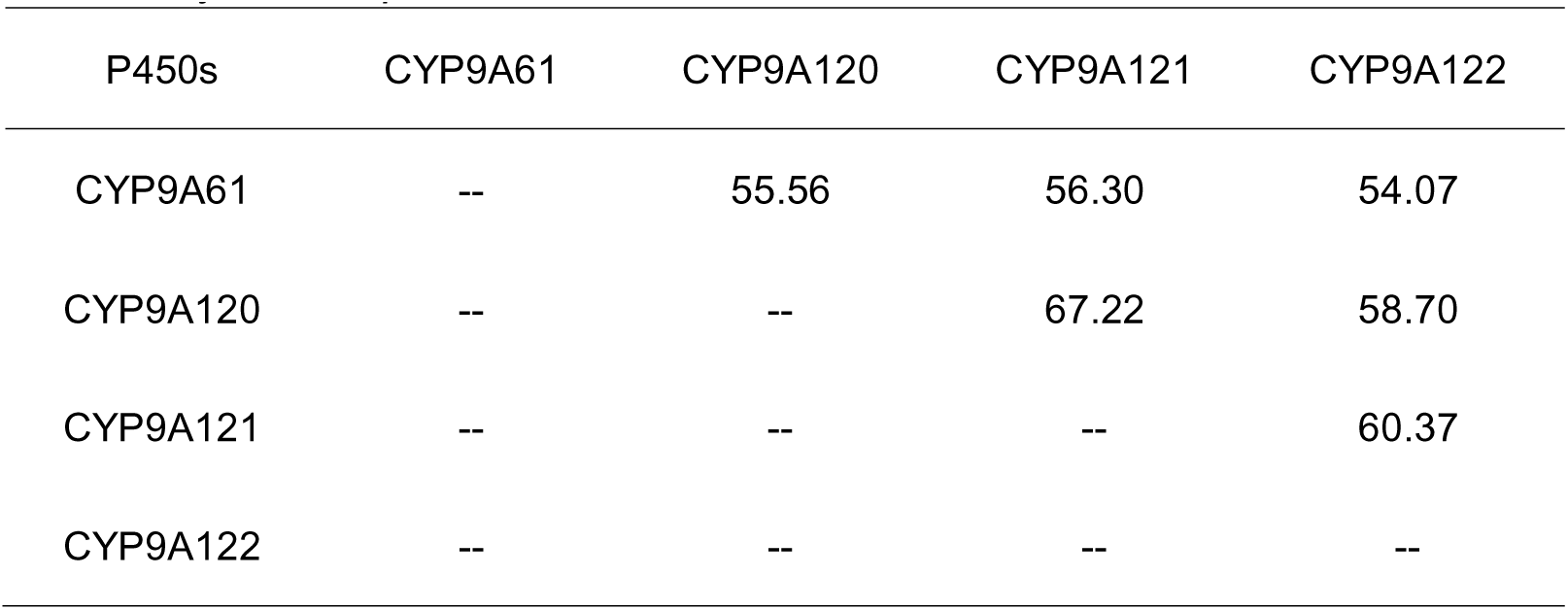
Identity (%) in amino acid sequence among P450s in CYP9A subfamily from *C.pomonella*.

**Table S3.** The number of CYP9A genes in different species of Lepidoptera.

| Species name | Number | References |
| --- | --- | --- |
| *Cydia pomonella* | 4 | 17 |
| *Chilo suppressalis* | 4 | Ma et al., 2019 |
| *Bombyx mori* | 4 | Lu et al., 2020 |
| *Helicoverpa armigera* | 6 | Pearce et al., 2017 |
| *Plutella xylostella* | 6 | You et al., 2013 |
| *Spodoptera litura* | 15 | 59 |
| *Spodoptera exigua* | 10 | 16 |
| *Spodoptera frugiperda* | 14 | Gouin et al., 2017; Xiao et al., 2020 |

---

**Table S4.** Information for P450s used for phylogenetic analysis.

| Insect species | P450s | GenBank No. |
| --- | --- | --- |
| *H. armigera* | CYP9A3 | KM016754.1 |
|  | CYP9A17 | AY753201.1.1 |
|  | CYP9A18 | KR095604.1 |
|  | CYP9A34 | KM016755.1 |
| *Spodoptera exigua* | CYP9A11 | KX443438.1 |
|  | CYP9A27 | KX443439.1 |
|  | CYP9A30 | MN179465.1 |
|  | CYP9A105 | KY348418.1 |
| *Spodoptera frugiperda* | CYP9A27 | MN480665.1 |
|  | CYP9A58 | MZ945714.1 |
|  | CYP9A59 | MZ945725.1 |
|  | CYP9A60 | MZ945685.1 |
| *Bombyx mori* | CYP9A19 | NM_001109934.1 |
|  | CYP9A20 | EF421989.1 |
|  | CYP9A21 | NM_001109924.1 |
|  | CYP9A22 | NM_001102464.2 |
| *Depressaria pastinacella* | CYP9A6 | AY295776.1 |
|  | CYP9A7 | AY295775.1 |
| *Chilo suppressalis* | CYP9A68 | KF701145.1 |

## Figure legends

**Fig. S1.**
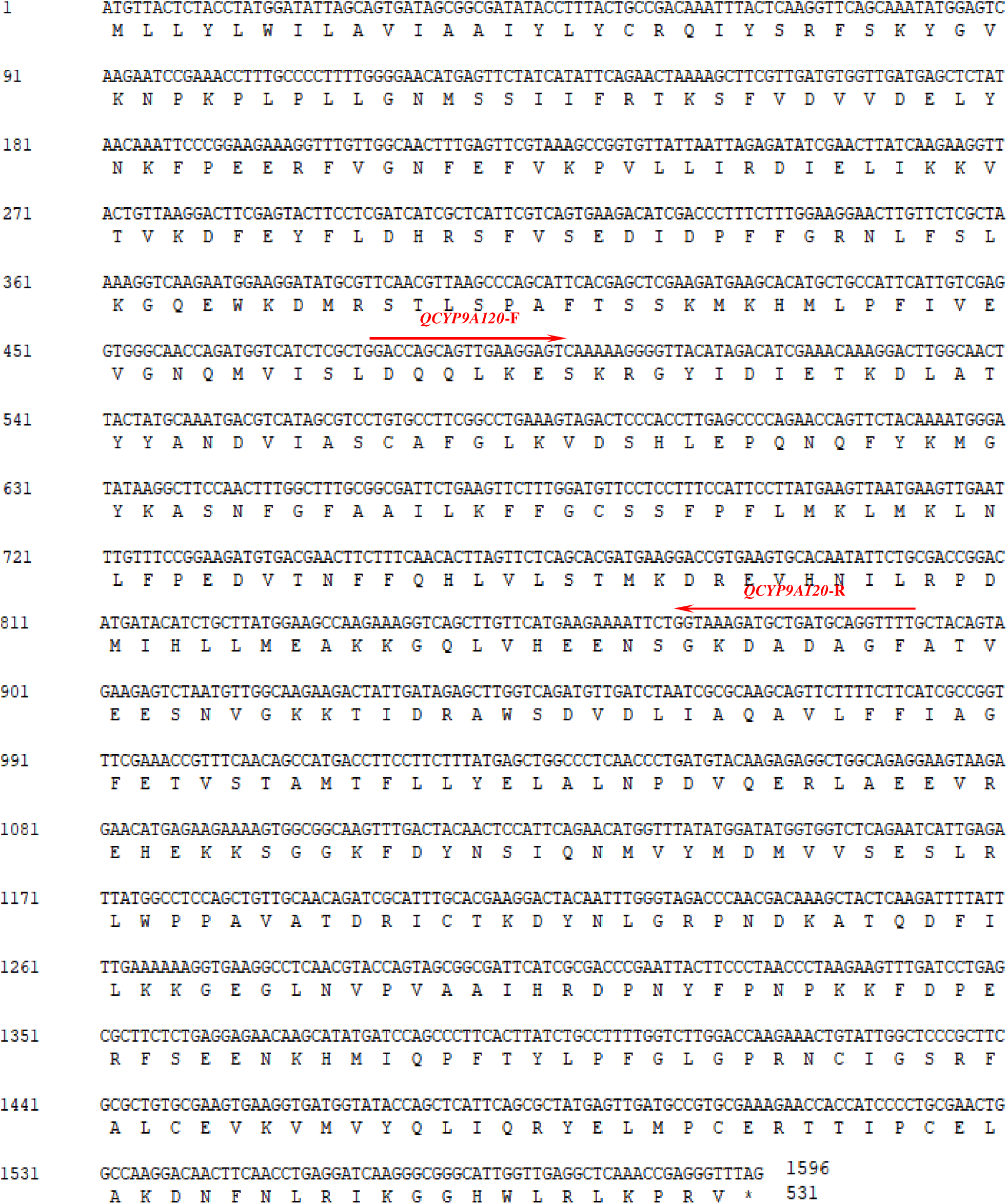
Nucleotide and deduced amino acid sequences of *CYP9A120* from *C. pomonella*. The primer used for RT-qPCR was marked with a red arrow.

**Fig. S2.**
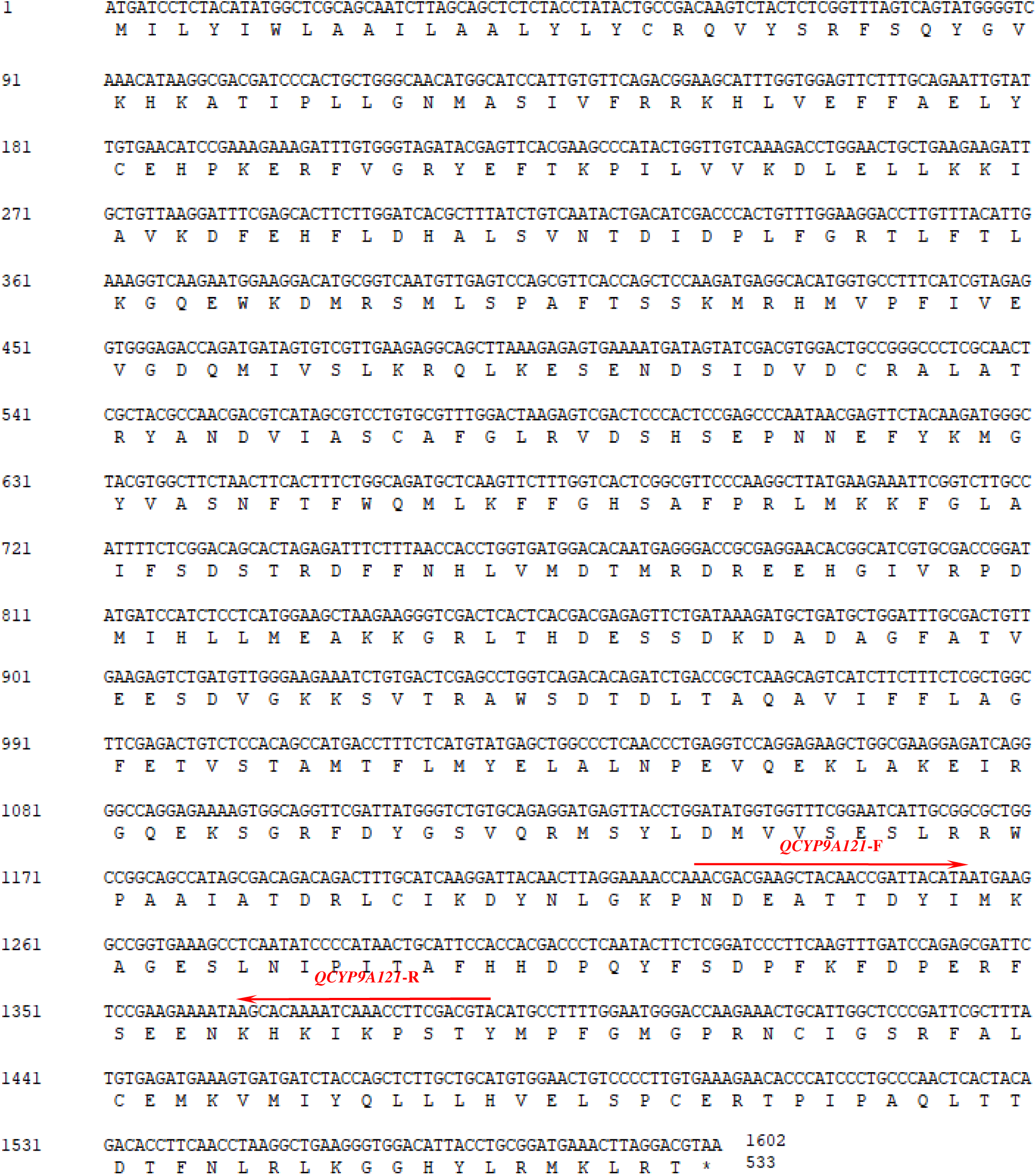
Nucleotide and deduced amino acid sequences of *CYP9A121* from *C. pomonella*. The primer used for RT-qPCR was marked with a red arrow.

**Fig. S3.**
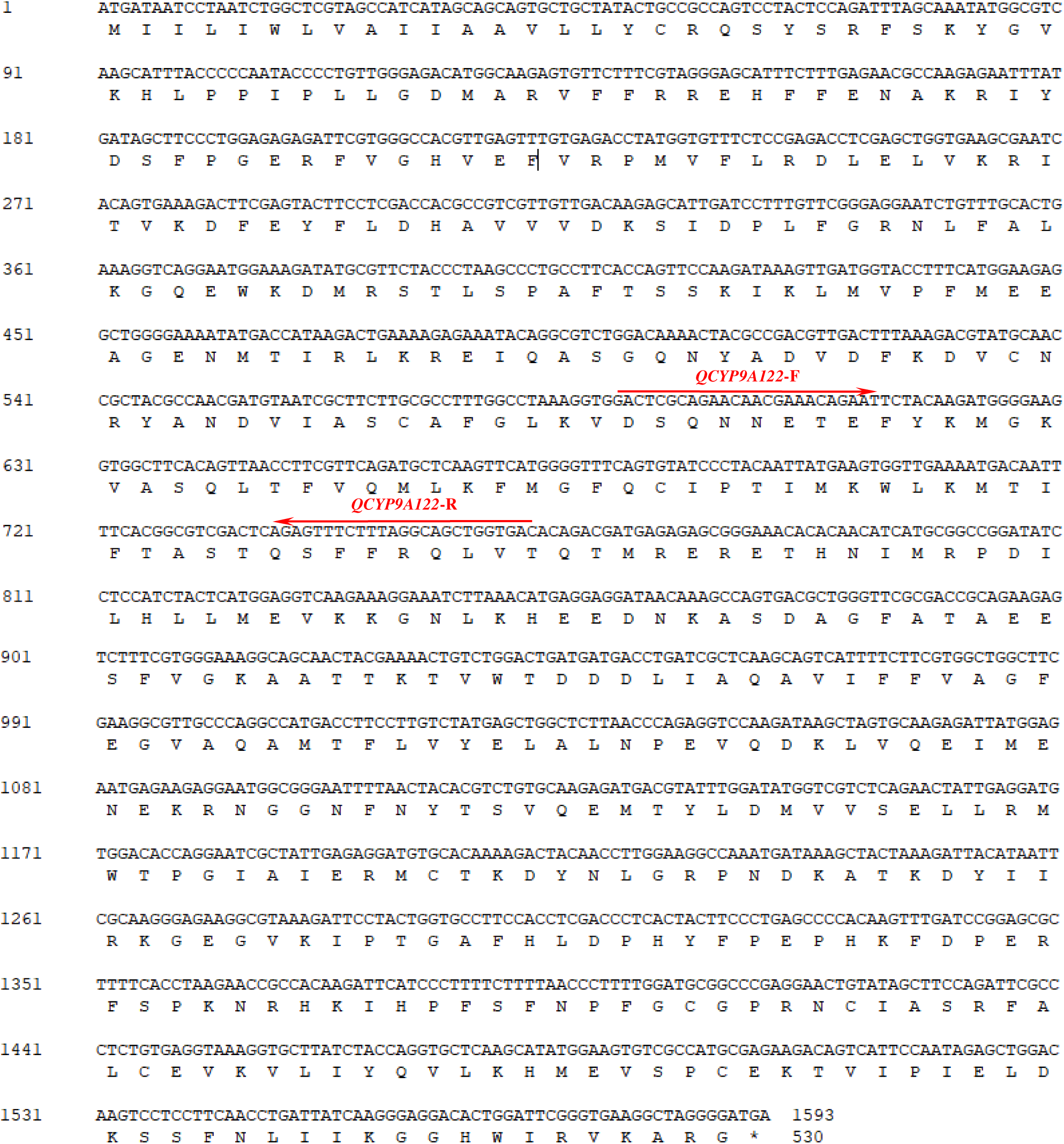
Nucleotide and deduced amino acid sequences of *CYP9A122* from *C. pomonella*. The primer used for RT-qPCR was marked with a red arrow.

**Fig. S4.**
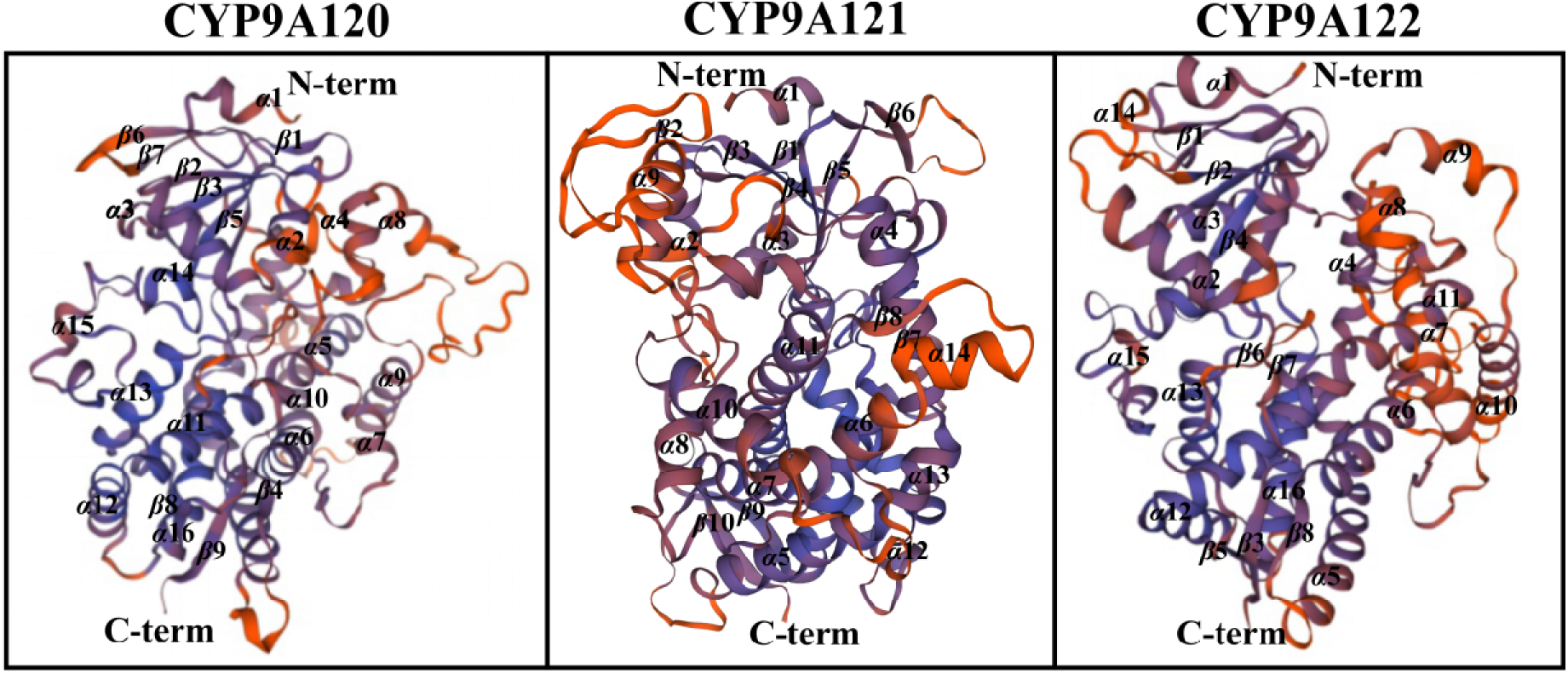
Predicted 3D structure of CYP9A120, CYP9A121, and CYP9A122 from *C. pomonella*. 3D structure of CYP9A120, CYP9A121 and CYP9A122 were predicted using homology modeling (automated mode) using the SWISS-MODEL (http://swissmodel.expasy.org). The target template sequence was searched using BLAST against the primary amino acid sequence contained in the SWISS-MODEL template library. A total of 50 templates were found. For each identified template, the template’s quality was predicted from features of the target-template alignment. The templates with the highest quality were then selected for model building. Thus, a 1.93 Å human cytochrome P450 3A4 (PDB no. 4d6z.1.A) bound to imidazole and an inhibitor were selected and used as the templates for CYP9A120 and CYP9A121, while a 2.15 Å human fetal-specific CYP3A7 (PDB no. 7mk8.1.A) bound to dithiothreitol was selected and used as the templates for CYP9A122. The N-term, C-term, and the α-helix and β-fold of each P450 were marked.

**Fig. S5.**
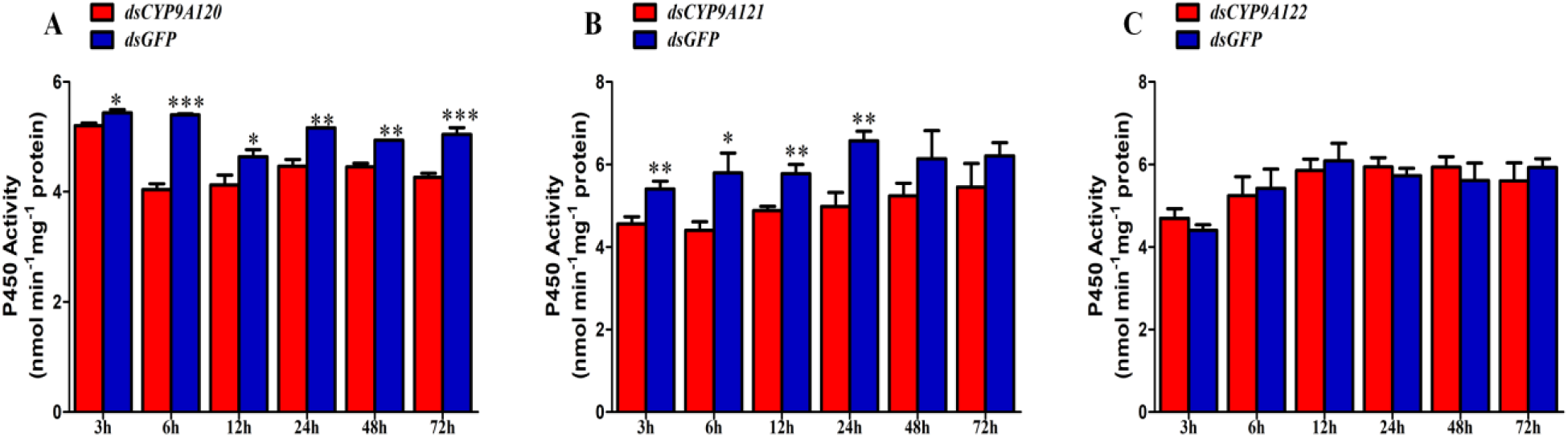
Effects of silencing of *CYP9A* genes on P450 enzyme activity of *C. pomonella*. Asterisks on the error bar indicate significant differences analyzed by the one-way analysis of variance (ANOVA) with the Student’s *t*-test (\**P* < 0.05, ** *P* < 0.01, *** *P* < 0.001).

**Fig. S6.**
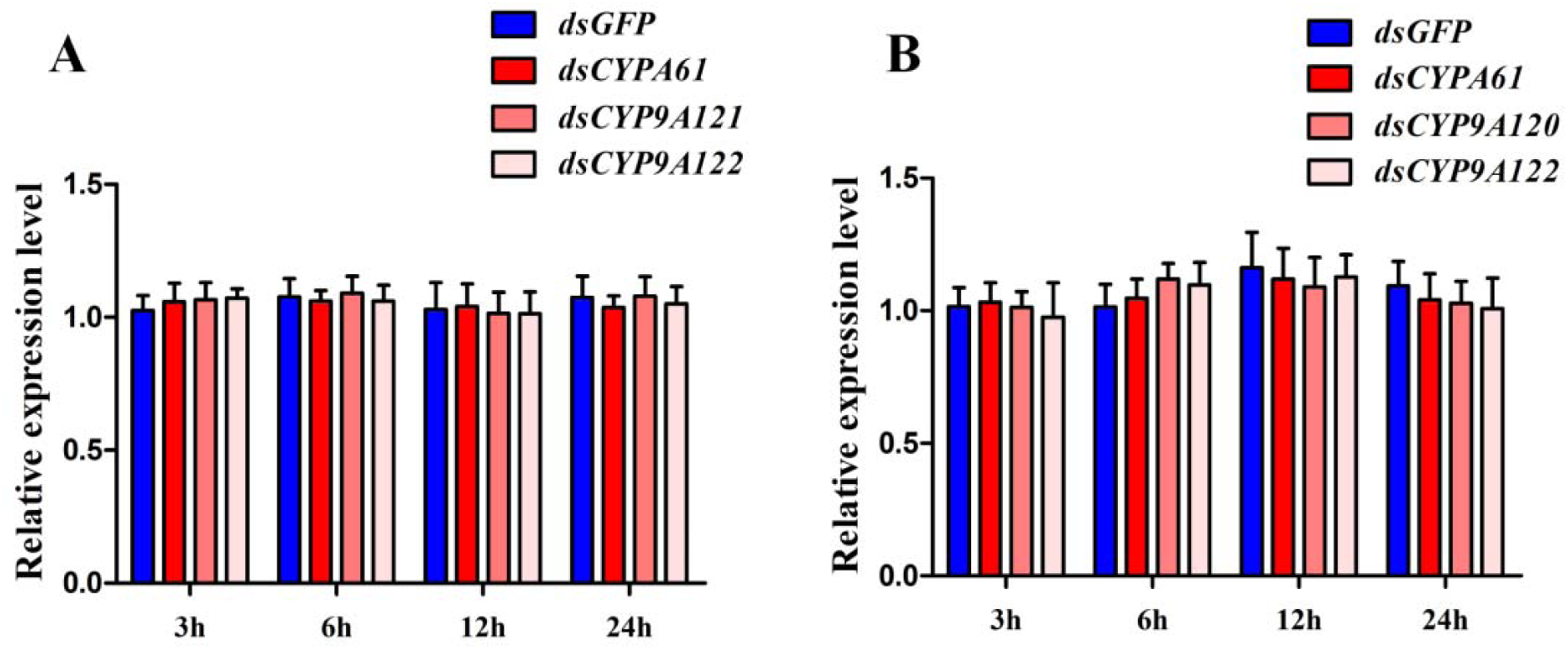
Detection of off-target effects after silencing of *CYP9A120* and *CYP9A121*. The results are shown as the mean ± SD.

**Fig. S7.**
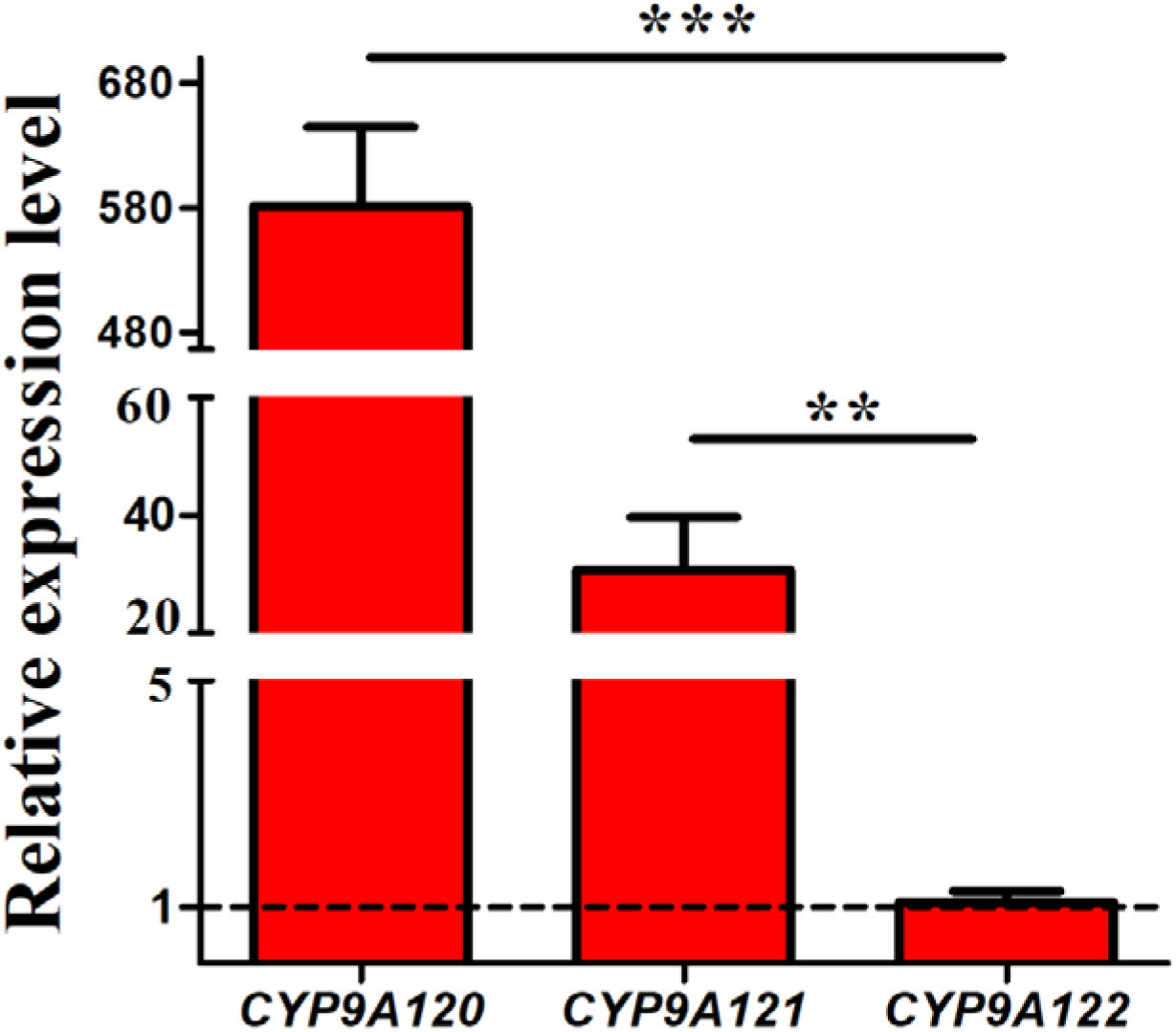
Expression profiles of *CYP9A* genes in third-instar larvae of *C. pomonella*. The results are shown as the mean ± SD. The error bars represent the standard errors calculated from three replicates. Asterisks on the error bar indicate significant differences analyzed by the one-way analysis of variance (ANOVA) with the Student’s *t*-test (** *P* < 0.01, *** *P* < 0.001).

## Notes

### Competing Interest Statement

The authors have declared no competing interest.

## REFERENCES

1. Yang XQ and Zhang YL, Investigation of insecticide-resistance status of *Cydia pomonella* in Chinese populations. B Entomol Res 105: 316–325 (2015).

2. Deligeorgidis NP, Kavallieratos NG, Malesios C, Sidiropoulos G, Deligeorgidis PN., Benelli G, Papanikolaou NE, Evaluation of combined treatment with mineral oil, fenoxycarb and chlorpyrifos against *Cydia pomonella*, *Phyllonorycter blancardella* and *Synanthedon myopaeformis* in apple orchards. Entomol Gen 39: 117–126 (2019).

3. Hu C, Wang W, Ju D, Chen G, Tan X, Mota-Sanchez D, Yang X, Functional characterization of a novel λ-cyhalothrin metabolising glutathione *S*-transferase, *CpGSTe3*, from the codling moth *Cydia pomonella*. Pest Manag Sci 76:1039–1047 (2019).

4. Wan FH, Yin CL, Tang R, Chen MH, Wu Q, Huang C, et al., A chromosome-level genome assembly of *Cydia pomonella* provides insights into chemical ecology and insecticide resistance. Nat Commun 10:4237(2019).

5. Wang W, Hu C, Li X, Wang X, Yang X, *CpGSTd3* is a *lambda*-Cyhalothrin metabolizing glutathione *S*-transferase from *Cydia pomonella* (L.). J Agric Food. Chem 67: 1165–1172 (2019).

6. Zhang X, Zhou, S, Wang Y, Chen Y, Research on Yining *Cydia pomonella* and other fruit insect investigations (2). Xinjiang. AgrSci 14 (1957).

7. Yang X, Wu Z, Zhang Y, Barros-Parada W, Toxicity of six insecticides on codling moth (Lepidoptera: Tortricidae) and effect on expression of detoxification genes. J Econ Entomol 109: 320–326 (2016).

8. Ju D, Mota-Sanchez D, Fuentes-Contreras E, Zhang Y, Wang X, Yang X, Insecticide resistance in the *Cydia pomonella* (L): Global status, mechanisms, and research directions. Pestic Biochem Physiol 178: 104925 (2021).

9. Reyes M, Franck P, Charmillot PJ, Ioriatti C, Olivares J, Pasqualini E, Sauphanor B, Diversity of insecticide resistance mechanisms and spectrum in European populations of the codling moth, *Cydia pomonella*. Pest Manag Sci 63: 890–902 (2007).

10. Hu C, Liu J, Wang W, Mota-Sanchez D, He S, Shi Y, Yang X, Glutathione *S*-Transferase genes are involved in *lambda*-cyhalothrin resistance in *Cydia pomonella* via sequestration. J Agric Food Chem 70: 2265–2279 (2022).

11. Desneux N, Decourtye A, Delpuech J, The sublethal effects of pesticides on beneficial arthropods. Annu Rev Entomol 52: 81–106 (2007).

12. Wang X, Xu X, Ullah F, Ding Q, Gao X, Desneux N, Song D, Comparison of full-length transcriptomes of different imidacloprid-resistant strains of *Rhopalosiphum padi* (Linné). Entomol Gen 41: 289–304 (2021).

13. Goff G.L, Nauen R, Recent Advances in the Understanding of Molecular Mechanisms of Resistance in Noctuid Pests. Insects 12: 674 (2021).

14. Nelson DR, Cytochrome P450 diversity in the tree of life. Biochim Biophys Acta Proteins Proteom 1866: 141–154 (2018).

15. Nauen R, Bass C, Feyereisen R, Vontas J, The Role of Cytochrome P450s in insect toxicology and resistance. Annu Rev Entomol 67: 105–124 (2022).

16. Shi Y, Jiang Q, Yang Y, Feyereisen R, Wu Y, Pyrethroid metabolism by eleven *Helicoverpa armigera* P450s from the CYP6B and CYP9A subfamilies. Insect Biochem Mol Biol 135: 103597 (2021).

17. Yang X, Li X, Zhang Y, 2013. Molecular cloning and expression of *CYP9A61*: a chlorpyrifos-ethyl and *lambda*-cyhalothrin-inducible cytochrome P450 cDNA from *Cydia pomonella*. Int J Mol Sci 14: 24211–24229 (2013).

18. Wei Z, Liu M, Hu C, Yang X, Overexpression of glutathione *S*-transferase genes in field λ-cyhalothrin-resistant population of *Cydia pomonella*: reference gene selection and expression analysis. J Agric Food Chem 68: 5825–5834 (2020).

19. Camacho C, Coulouris G, Avagyan V, Ma N, Papadopoulos J, Bealer K, Madden TL, BLAST+: architecture and applications. BMC Bioinformatics 10: 421 (2009).

20. Larkin MA, Blackshields G, Brown NP, Chenna R, McGettigan PA, McWilliam H, Valentin F, Wallace IM, Wilm A, Lopez R, Thompson JD, Gibson TJ, Higgins DG, Clustal W and Clustal X version 2.0. Bioinformatics 23: 2947–2948 (2007).

21. McGuffin LJ, Bryson K, Jones DT, The PSIPRED protein structure prediction server. Bioinformatics 16: 404–405 (2020).

22. Kumar S, Stecher G, Tamura K, MEGA7: molecular evolutionary genetics analysis version 7.0 for bigger datasets. Mol Biol Evol 33: 1870–1874 (2016).

23. Bustin SA, Benes V, Garson JA., Hellemans J, Huggett J, Kubista M, Mueller R, Nolan T, Pfaffl MW, Shipley,GL, Vandesompele J, Wittwer CT, The MIQE guidelines: minimum information for publication of quantitative real-time PCR experiments. Clin Chem 55: 611–622 (2009).

24. Livak KJ, Schmittgen TD, Analysis of relative gene expression data using real-time quantitative PCR and the 2^−ΔΔ^CT method. Methods 25: 402–408 (2001).

25. Bradford MM, A rapid and sensitive method for the quantitation of microgram quantities of protein utilizing the principle of protein-dye binding. Anal Biochem 72: 248–254 (1976).

26. Omura T, Sato R, The carbon monoxide-binding pigment of liver microsomes. I. evidence for its hemoprotein nature. J Biol Chem 239: 2370–2378 (1964).

27. Shi Y, Wang H, Liu Z, Wu S, Yang Y, Feyereisen R, Heckel DG, Wu Y, Phylogenetic and functional characterization of ten P450 genes from the CYP6AE subfamily of *Helicoverpa armigera* involved in xenobiotic metabolism. Insect Biochem Mol Biol 93: 79–91 (2018).

28. 28 Feyereisen R, Insect CYP genes and P450 enzymes. In: Gilbert, L.I. (Ed.), Insect Molecular Biology and Biochemistry. Elsevier B.V., London, pp. 236–316 (2012).

29. Brattsten LB, Holyoke CWJr, Leeper JR, Raffa KF, Insecticide resistance: challenge to pest management and basic research. Science 231:1255–1260 (1986).

30. Abbas N, Khan HAA, Shad SA, Resistance of the House Fly *Musca domestica* (Diptera: Muscidae) to Lambda-Cyhalothrin: Mode of Inheritance, Realized Heritability, and Cross-Resistance to Other Insecticides. Ecotoxicology 23: 791–801 (2014).

31. Guedes RNC, Walse SS, Throne JE, Sublethal exposure, insecticide resistance, and community stress. Curr Opin Insect Sci 21: 47–53 (2017).

32. Santoiemma G, Tonina L, Marini L, Duso C, Integrated management of *Drosophila suzukii* in sweet cherry orchards. Entomol Gen 40: 297–305 (2020).

33. Paula DP, Lozano RE, Menger JP, Andow D, Koch, RL, Identification of point mutations related to pyrethroid resistance in voltage-gated sodium channel genes in *Aphis glycines*. Entomol Gen 41: 243–255 (2021).

34. Li X, Schuler MA, Berenbaum MR, Molecular mechanisms of metabolic resistance to synthetic and natural xenobiotics. Annu Rev Entomol 52: 231–253 (2007).

35. Gul H, Ullah F, Biondi A, Desneux N, Qian D, Gao XW, Song DL, Resistance against clothianidin and associated fitness costs in the chive maggot, *Bradysia odoriphaga*. Entomol Gen 39: 81–92 (2019).

36. Saeed R, Abbas N, Hafez AM, Biological fitness costs in emamectin benzoate-resistant strains of *Dysdercus koenigii*. Entomol Gen 41: 267–278 (2021).

37. Shan JQ, Zhu B, Gu SH, Liang P, Gao XW, Development of resistance to chlorantraniliprole represses sex pheromone responses in male *Plutella xylostella*. Entomol Gen 41: 615–625 (2021).

38. Wang Z, Jiang S, Mota-Sanchez, D, Wang W, Li X, Gao Y, Lu X, Yang X, Cytochrome P450-mediated λ-cyhalothrin-resistance in a field strain of *Helicoverpa armigera* from northeast China. J Agric Food Chem 67: 3546–3553 (2019).

39. Yang X, Li X, Cang X, Guo J, Shen X, Wu K, Influence of seasonal migration on the development of the insecticide resistance of oriental armyworm (*Mythimna separata*) to λ-cyhalothrin. Pest Manag Sci 78: 1194–1205 (2022).

40. Yang X, Li X, Zhang Y, Molecular cloning and expression of *CYP9A61*: a chlorpyrifos-ethyl and *lambda*-cyhalothrin-inducible cytochrome P450 cDNA from *Cydia pomonella*. Int J Mol Sci 14: 24211–24229 (2013).

41. Feyereisen R, Arthropod CYPomes illustrate the tempo and mode in P450 evolution. Biochim Biophys Acta 1814: 19–28 (2011).

42. Berenbaum MR, Johnson RM, Xenobiotic detoxification pathways in honey bees. Curr Opin Insect Sci 10: 51–58 (2015).

43. Zhao X, Xu X, Wang X, Yin Y, Li M, Wu Y, Liu Y, Cheng Q, Gong C, Shen L, Mechanisms for multiple resistances in field populations of rice stem borer, *Chilo suppressalis* (Lepidoptera: Crambidae) from Sichuan Province, China. Pestic Biochem Physiol 171: 104720 (2021).

44. Li F, Ni M, Zhang H, Wang B, Xu K, Tian J, Hu J, Shen W, Li B, Expression profile analysis of silkworm P450 family genes after phoxim induction. Pestic Biochem Physiol 122: 103–109 (2015).

45. Huang Y, Shen G, Jiang H, Jiang X, Dou W, Wang J, Multiple P450 genes: Identification, tissue-specific expression and their responses to insecticide treatments in the oriental fruit fly, *Bactrocera dorsalis* (Hendel) (Diptera: Tephritidea). Pestic Biochem Physiol 106: 1–7 (2013).

46. Xu L, Zhao J, Sun Y, Xu D, Xu G, Xu X, Zhang Y, Huang S, Han Z, Gu Z, Constitutive overexpression of cytochrome P450 monooxygenase genes contribute to chlorantraniliprole resistance in *Chilo suppressalis* (Walker). Pest Manag Sci 75: 718–725 (2019).

47. Komagata O, Kasai S, Tomita T, Overexpression of cytochrome P450 genes in pyrethroid-resistant *Culex quinquefasciatus*. Insect Biochem Mol Biol 40: 146–152 (2010).

48. Zhou X, Ma C, Li M, Sheng C, Liu H, Qiu X, *CYP9A12* and *CYP9A17* in the cotton bollworm, *Helicoverpa armigera*: sequence similarity, expression profile and xenobiotic response. Pest Manag Sci 66: 65–73 (2010).

49. Lu K, Song Y, Zeng R, The role of cytochrome P450-mediated detoxification in insect adaptation to xenobiotics. Curr Opin Insect Sci 43: 103–107 (2021).

50. Amezian D, Nauen R, Goff GL, Transcriptional regulation of xenobiotic detoxification genes in insects-an overview. Pestic Biochem Physiol 174: 104822 (2021).

51. Vandenhole M, Dermauw W, Leeuwen TV, Short term transcriptional responses of P450s to phytochemicals in insects and mites. Curr Opin Insect Sci 43: 117–127 (2021).

52. Giraudo M, Hilliou F, Fricaux T, Audant P, Feyereisen R, Goff GL, Cytochrome P450s from the fall armyworm (*Spodoptera frugiperda*): responses to plant allelochemicals and pesticides. Insect Mol Biol 24: 115–128 (2015).

53. Wang R, Staehelin C, Xia Q, Su Y, Zeng R, Identification and characterization of *CYP9A40* from the tobacco cutworm moth (*Spodoptera litura*), a cytochrome P450 gene induced by plant allelochemicals and insecticides. Int J Mol Sci 16: 22606–22620 (2015).

54. Zhu W, Yu R, Wu H, Zhang X, Liu Y, Zhu K, Zhang J, Ma E, Identification and characterization of two CYP9A genes associated with pyrethroid detoxification in *Locusta migratoria*. Pestic Biochem Physiol 132: 65–71 (2016).

55. Rose RL, Goh D, Thompson DM, Verma KD, Heckel DG, Gahan LJ, Roe RM, Hodgson E, Cytochrome P450 *(CYP)9A1* in *Heliothis virescens*: the first member of a new CYP family. Insect Biochem Mol Biol 27: 0–615 (1997).

56. Hilliou F, Chertemps T, Maïbèche M, Goff GL, Resistance in the genus *Spodoptera*: key insect detoxification genes. Insects 12: 544 (2021).

57. Xu L, Li D, Qin J, Zhao W, Qiu L, Over-expression of multiple cytochrome P450 genes in fenvalerate-resistant field strains of *Helicoverpa armigera* from north of China. Pestic Biochem Physiol 132: 53–58 (2016).

58. Vandenhole M, Dermauw W, Leeuwen TV, Short term transcriptional responses of P450s to phytochemicals in insects and mites. Curr Opin Insect Sci 43: 117–127 (2021).

59. Cheng T, Wu J, Wu Y, Chilukuri RV, Huang L, Yamamoto K, Feng L, Li W, Chen Z, Guo H, Liu J, Li S, Wang X, Peng L, Liu D, Guo Y, Fu B, Li Z, Liu C, Chen Y, Tomar A, Hilliou F, Montagné N, Jacquin-Joly E, d’Alençon E, Seth RK, Bhatnagar RK, Jouraku A, Shiotsuki T, Kadono-Okuda K, Promboon A, Smagghe G, Arunkumar KP, Kishino H, Goldsmith MR, Feng Q, Xia Q, Mita K, Genomic adaptation to polyphagy and insecticides in a major East Asian noctuid pest. Nat Ecol Evol 1: 1747–1756 (2017).

60. Wang R, Liu S, Baerson SR, Qin Z, Ma Z, Su Y, Zhang J, Identification and functional analysis of a novel cytochrome P450 gene *CYP9A105* associated with pyrethroid detoxification in *Spodoptera exigua* Hübner. Int J Mol Sci 19: 737 (2018).

61. Terenius O, Papanicolaou A, Garbutt JS, Eleftherianos I, Huvenne H, et al., RNA interference in Lepidoptera: an overview of successful and unsuccessful studies and implications for experimental design. J Insect Physiol 57: 231–245 (2011).

62. Cooper AM, Silver K, Zhang J, Park Y, Zhu KY, Molecular mechanisms influencing efficiency of RNA interference in insects. Pest Manag Sci 75: 18–28 (2019).

63. Yang Y, Yue L, Chen S, Wu Y, Functional expression of *Helicoverpa armigera* CYP9A12 and CYP9A14 in *Saccharomyces cerevisiae*. Pestic Biochem Physiol 92: 101–105 (2008).

64. Itokawa K, Komagata O, Kasai S, Okamura Y, Masada M, Tomita T, Genomic structures of *Cyp9m10* in pyrethroid resistant and susceptible strains of *Culex quinquefasciatus*. Insect Biochem Mol Biol 40: 631–640 (2010).

65. Schmidt JM., Good RT., Appleton B, Sherrard J, Raymant GC, Bogwitz MR, Martin J, Daborn PJ, Goddard ME, Batterham P, Robin C, Copy number variation and transposable elements feature in recent, ongoing adaptation at the *Cyp6g1* locus. PLOS Genet 6: e1000998 (2010).

66. Bass C, Zimmer CT, Riveron JM, Wilding CS, Wondji CS, Kaussmann M, Field LM, Williamson MS, Nauen R, Gene amplification and microsatellite polymorphism underlie a recent insect host shift. PNAS 110: 19460–19465 (2013).

67. Zimmer CT, Garrood WT, Singh KS, Randall E, Lueke B, Gutbrod O, Matthiesen S, Kohler M, Nauen R, Davies TGE, Bass C, Neofunctionalization of duplicated P450 genes drives the evolution of insecticide resistance in the brown planthopper. Curr Biol 28: 268–274 (2018).

68. Gimenez S, Abdelgaffar H, Goff GL, Hilliou F, Blanco CA, Hännigerm S, Bretaudeau A, Legeai F, Nègre N, Jurat-Fuentes JL, d’Alençon E, Nam K, Adaptation by copy number variation increases insecticide resistance in the fall armyworm. Commun Biol 3: 664 (2020).

## Reference

Cheng T, Wu J, Wu Y, Chilukuri RV, Huang L, Yamamoto K, Feng L, Li W, Chen Z, Guo H, Liu J, Li S, Wang X, Peng L, Liu D, Guo Y, Fu B, Li Z, Liu C, Chen Y, Tomar A, Hilliou F, Montagné N, Jacquin-Joly E, D Alençon E, Seth R.K, Bhatnagar RK, Jouraku A, Shiotsuki T, Kadono-Okuda K, Promboon A, Smagghe G, Arunkumar KP, Kishino H, Goldsmith MR, Feng Q, Xia Q, Mita K. Genomic adaptation to polyphagy and insecticides in a major East Asian noctuid pest. Nat Ecol Evol 1: 1747–1756 (2017).

Gouin A, Bretaudeau A, Nam K, Gimenez S, Aury J, Duvic B, Hilliou F, Durand N, Montagné N, Darboux I, Kuwar S, Chertemps T, Siaussat D, Bretschneider A, Moné Y, Ahn S, Hänniger S, Grenet AG, Neunemann D, Maumus F, Luyten I, Labadie K, Xu W, Koutroumpa F, Escoubas J, Llopis A, Maïbè che-Coisne M, Salasc F, Tomar A, Anderson AR, Khan SA, Dumas P, Orsucci M, Guy J, Belser C, Alberti A, Noel B, Couloux A, Mercier J, Nidelet S, Dubois E, Liu N, Boulogne I, Mirabeau O, Le Goff G, Gordon K, Oakeshott J, Consoli FL, Volkoff A, Fescemyer HW, Marden JH, Luthe DS, Herrero S, Heckel DG, Wincker P, Kergoat GJ, Amselem J, Quesneville H, Groot AT, Jacquin-Joly E, Nègre N, Lemaitre C, Legeai F, D Alençon E, Fournier P. Two genomes of highly polyphagous lepidopteran pests (*Spodoptera frugiperda*, Noctuidae) with different host-plant ranges. Sci Rep 7: 11816 (2017).

Lu F, Wei Z, Luo Y, Guo H, Zhang G, Xia Q, Wang Y. SilkDB 3.0: Visualizing and exploring multiple levels of data for silkworm. Nucleic Acids Res 48: D749–D755 (2020).

Ma W, Zhao X, Yin C, Fan J, Du X, Chen T, Zhang Q, Qiu L, Xu H, Hull, J J, Li G, Sung W, Li F, Lin Y. A chromosome-level genome assembly reveals the genetic basis of cold tolerance in a notorious rice insect pest *Chilo suppressalis*. Mol Ecol Resour 20: 268–282 (2019).

Pearce SL, Clarke DF, East PD, Elfekih S, Gordon KHJ, Jermiin LS, McGaughran A, Oakeshott JG, Papanikolaou A, Perera OP, Rane RV, Richards S, Tay WT, Walsh TK, Anderson A, Anderson CJ, Asgari S, Board PG, Bretschneider A, Campbell PM, Chertemps T, Christeller JT, Coppin CW, Downes SJ, Duan G, Farnsworth CA, Good RT, Han LB, Han YC, Hatje K, Horne I, Huang YP, Hughes DST, Jacquin-Joly E, James W, Jhangiani S, Kollmar M, Kuwar SS, Li S, Liu N, Maibeche MT, Miller JR, Montagne N, Perry T, Qu J, Song SV, Sutton GG, Vogel H, Walenz BP, Xu W, Zhang H, Zou Z, Batterham P, Edwards OR, Feyereisen R, Gibbs RA, Heckel DG, McGrath A, Robin C, Scherer SE, Worley KC, Wu Y.D. Genomic innovations, transcriptional plasticity and gene loss underlying the evolution and divergence of two highly polyphagous and invasive *Helicoverpa* pest species. BMC Biol 15: 63 (2017).

Xiao H, Ye X, Xu H, Mei Y, Yang Y, Chen X, Yang Y, Liu T, Yu Y, Yang W, Lu Z, Li F. The genetic adaptations of fall armyworm *Spodoptera frugiperda* facilitated its rapid global dispersal and invasion. Mol Ecol Resour 20: 1050–1068 (2020).

You M, Yue Z, He W, Yang X, Yang G, Xie M, Zhan D, Baxter SW, Vasseur L, Gurr GM, Douglas CJ, Bai J, Wang P, Cui K, Huang S, Li X, Zhou Q, Wu Z, Chen Q, Liu C, Wang B, Li X, Xu X, Lu C, Hu M, Davey JW, Smith SM, Chen M, Xia X, Tang W, Ke F, Zheng D, Hu Y, Song F, You Y, Ma X, Peng L, Zheng Y, Liang Y, Chen Y, Yu L, Zhang Y, Liu Y, Li G, Fang L, Li J, Zhou X, Luo Y, Gou C, Wang J, Wang J, Yang H, Wang J. A heterozygous moth genome provides insights into herbivory and detoxification. Nat Genet 45: 220–225 (2013).

